# Environmental stress and phenotypic tradeoff modulate the adaptive potential of novel coding sequences for *de novo* gene birth

**DOI:** 10.64898/2026.08.24.746591

**Authors:** Lin Chou, Nelson Castilho Coelho, Saurin Parikh, Catherine Douds, John Iannotta, Aaron Wacholder, Jiwon Lee, Carly Houghton, Anne-Ruxandra Carvunis

**Affiliations:** Department of Computational and Systems Biology, School of Medicine, University of Pittsburgh, Pittsburgh, PA 15213, USA; Pittsburgh Center for Evolutionary Biology and Medicine, School of Medicine, University of Pittsburgh, Pittsburgh, PA 15213, USA; Institute of Molecular Systems Biology, Department of Biology, ETH Zurich, Zurich, Switzerland

## Abstract

Novel protein-coding genes can emerge *de novo* from ancestrally noncoding sequences and promote adaptation to environmental stresses. Previous work proposes that pervasive translation of lowly expressed open reading frames (ORFs) in noncoding regions creates a rich reservoir of “proto-genes,” of which subsequent acquisition of gene-like properties, such as increased expression, may be favored or purged by natural selection depending on their phenotypic impact. However, whether and how environmental conditions affect the phenotypic impact of proto-genes remains unclear. Here, we experimentally simulated proto-gene evolution in *Saccharomyces cerevisiae* by individually increasing the expression of nearly a thousand *de novo* ORFs with prior evidence of native translation under osmotic and endoplasmic-reticulum stress and in control environments. High-throughput phenotyping revealed that growth effects of increased expression varied strongly across environments for *de novo* ORFs. A follow-up screen across 22 diverse environments revealed a robust positive correlation between environmental stress severity and the mean growth effects of increased *de novo* ORF expression. At the individual level, 5.4% of tested *de novo* ORFs conferred beneficial phenotypes in at least one environment, and 83.3% of these also caused deleterious effects elsewhere, revealing widespread phenotypic tradeoffs. We demonstrate that increased expression of the *de novo* translated ORF YLR112W results in increased growth in the presence of rapamycin through general dampening of the growth-repressing transcriptomic response induced by this drug. Together, these findings demonstrate that stress severity shapes the phenotypic consequences of increased proto-gene expression and shed light on tradeoffs and transcriptome remodeling as mechanisms underlying such environmental dependency.

## Introduction

Proteins play a fundamental role in biological processes, making understanding the functions of translated sequences in genomes one of the major quests in biology. The quest has further expanded over the past decade, as ribosome-profiling (ribo-seq) studies have revealed pervasive translation of open reading frames (ORFs) in addition to annotated protein-coding genes (Ingolia, et al. 2009; Carvunis, et al. 2012; Ruiz-Orera and Albà 2019; Wacholder, et al. 2023). Most of these unannotated translated ORFs are short and have low expression levels (Sandmann, et al. 2023; Wacholder, et al. 2023). Although a small fraction of unannotated translated ORFs revealed by ribo-seq have been characterized in various biological contexts (Chen, et al. 2020; Prensner, et al. 2021; Sandmann, et al. 2023; Wacholder, et al. 2023; Zheng, et al. 2023; Hofman, et al. 2024), functional understanding of the majority is still lacking, as they had been overlooked before the recent advent of ribo-seq.

Evolutionary analyses have revealed that most of the unannotated translated ORFs have recent *de novo* evolutionary origins in noncoding sequences of recent ancestors (Carvunis, et al. 2012; Van Oss and Carvunis 2019; Sandmann, et al. 2023; Rich, et al. 2024; Vara, et al. 2024; Zhao, et al. 2024). The recent evolutionary origins make these sequences only present in narrow ranges of species or populations (Carvunis, et al. 2012; Van Oss and Carvunis 2019; Sandmann, et al. 2023; Rich, et al. 2024; Vara, et al. 2024; Zhao, et al. 2024). This property poses a major challenge in the functional understanding of these sequences: it makes in silico homology-based prediction of protein function or domain unfruitful, emphasizing the need for direct experimental testing (Van Oss and Carvunis 2019; Zhao, et al. 2024). The unannotated translatome and the *de novo* translatome largely overlap (Sandmann, et al. 2023; Wacholder, et al. 2023), so the functional investigation of these two groups is synergistic.

*De novo* translated ORFs are often thought of as a source for new protein-coding genes and adaptation (Van Oss and Carvunis 2019; Ardern and uz-Zaman 2023; Zhao, et al. 2024). One evolutionary model representing this train of thought is the proto-gene model of *de novo* gene birth, which proposes that *de novo* translated ORFs, or proto-genes, may gain “gene-like” properties, such as longer sequences or higher expression, during evolution and undergo natural selection, leading them either toward gene birth or return to noncoding status (Carvunis, et al. 2012). Supporting this model, several studies have reported *de novo* emergence of translated ORFs from ancestrally noncoding regions and the involvement of these translated sequences in various lineage-specific traits or adaptation to stress environment (Baalsrud, et al. 2018; Van Oss and Carvunis 2019; Zhuang, et al. 2019; Weisman 2022; Wacholder, et al. 2023; uz-Zaman, et al. 2024; Zhao, et al. 2024; Jin, et al. 2025; Li, et al. 2025). The possibility of future refinement of *de novo* translated ORFs towards organismal adaptation emphasizes the need to expand the scope of functional investigation of *de novo* translated ORFs. Current physiological implications and potential function in the future upon acquisition of gene-like properties, or “adaptive potential”, are both important puzzles to understand the biological significance of these sequences.

High-throughput genetic screens of unannotated translated ORFs or *de novo* sequences have provided fruitful results for functional understanding of these sequences. Knockout or knockdown experiments using CRISPR or RNA interference have discovered unannotated translated ORFs involved in the growth of human and yeast cells (Chen, et al. 2020; Prensner, et al. 2021; Wacholder, et al. 2023; Zheng, et al. 2023; Hofman, et al. 2024; Pai, et al. 2025; Schlesinger, et al. 2025; Valdivia-Francia, et al. 2025) and *de novo* sequences involved in reproduction in fruit flies (Reinhardt, et al. 2013; Rivard, et al. 2021). How acquisition of gene-like properties affects phenotypes is relatively understudied. One study used overexpression as a strategy to gain functional insight into the potential evolutionary changes to *de novo* ORFs, increased expression in this case, and has discovered several sequences for which overexpression provides beneficial phenotypes (Vakirlis, et al. 2020). However, limitations exist in these prior successes: these functional studies are biased toward human cell lines, have a limited number of sequences, have tested too few environmental contexts to gain a fuller picture of functionality, and are mostly focused on gene deletion rather than testing their potential for gene birth.

In this study, we ask what the phenotypic impact of increased *de novo* translated ORFs expression is over diverse environments using yeast *Saccharomyces cerevisiae* as a model. More specifically, how does environmental stress shift the phenotypic impact of increased *de novo* translated ORF expression? Are phenotypes resulting from increased expression of *de novo* translated ORFs environment-specific? And what are the molecular mechanisms enabling a *de novo* translated ORF to confer an environment-specific beneficial phenotype? Our aim is to expand the proto-gene model of *de novo* gene birth and explicitly model how the phenotypic impact of increased proto-gene expression depends on the environment. Increased expression of individual proto-genes may have beneficial, deleterious, or no impact on phenotypes, which in turn dictates whether, and in which direction, selection may act on the increased expression **(Figure 1A)**. A quantitative estimate of how often each of these scenarios take place relative to each other, and a qualitative understanding of which environment may be more or less permissive to *de novo* gene birth, is needed.

**Figure 1.**
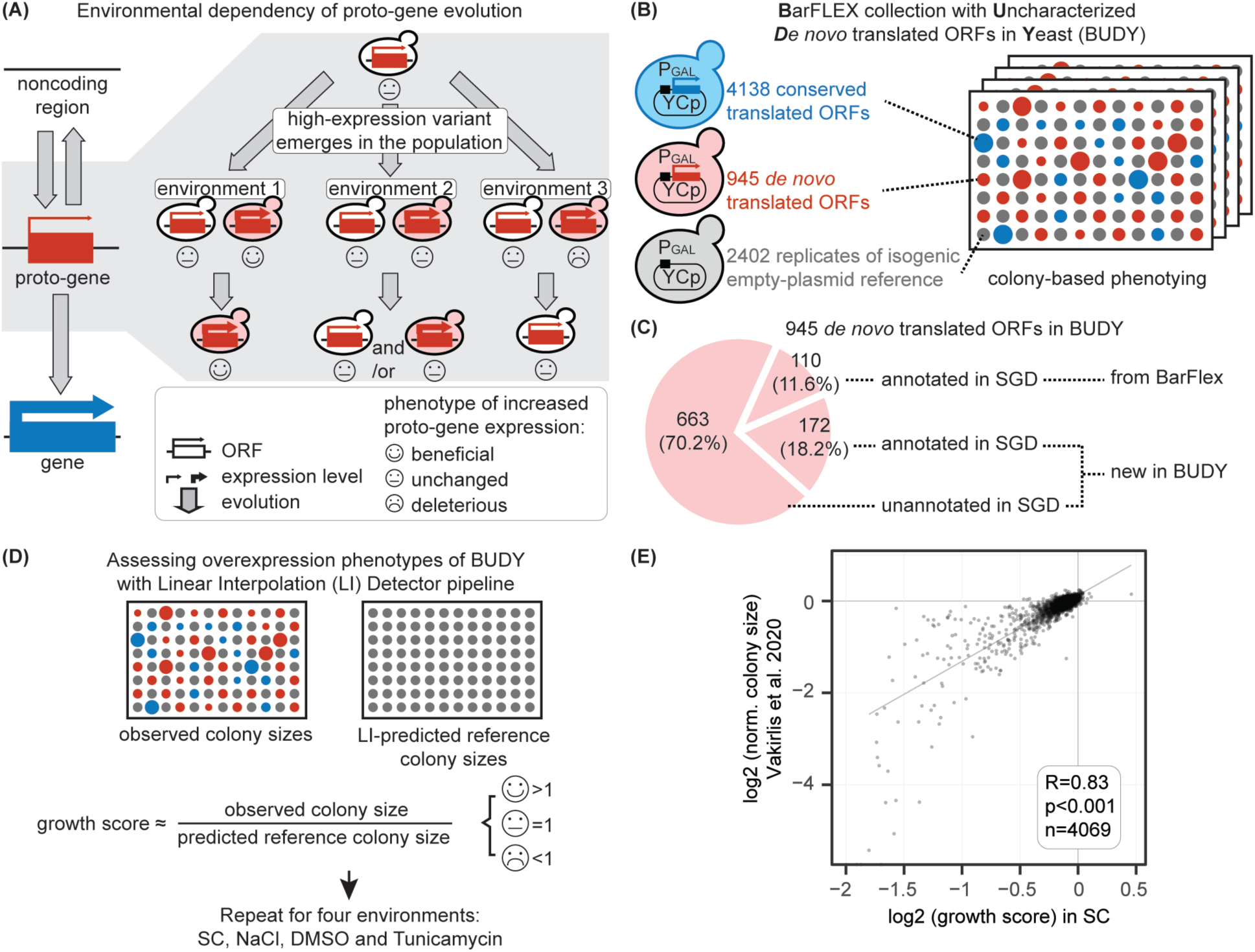
Environmental dependency of proto-gene evolution. (A) Theoretical model of the environmental dependency of proto-gene evolution. Protein-coding genes may emerge *de novo* from previously noncoding regions through an intermediary state called proto-genes (Carvunis, et al. 2012). Proto-genes are shorter and have lower expression levels compared to genes. Mutations that make proto-genes more “gene-like”, such as those that increase expression level (bold arrow), may occur during evolution. We ask whether increased proto-gene expression has a universal impact across environments, or environment-specific impact. Variants that have a deleterious phenotype might be selected against during evolution (environment 3), whereas variants with no (environment 2) or even beneficial (environment 1) effects may be retained during evolution through drift or adaptive evolution. If the phenotypic impact is environment-specific, the success of *de novo* gene birth is dependent on the succession of specific environments experienced by the organism over evolutionary time. (B)-(D) Assessing phenotypic impact of increased expression over different environments. (B) We experimentally simulated proto-gene evolution in four environments by individually overexpressing translated ORFs with recent *de novo* evolutionary origins within the *Saccharomyces* genus and known native translation in *Saccharomyces cerevisiae* using BUDY, an overexpression collection based on the BarFLEX collection (Douglas, et al. 2012). In total, the library includes overexpression strains of 945 *de novo* translated ORFs, 4,138 conserved translated ORFs, and 2,402 replicates of an isogenic empty plasmid reference strain as controls. Note that “proto-gene” is a concept, and we use *de novo* translated ORFs as an empirical proxy for proto-genes. (C) Composition of 945 *de novo* translated ORFs in BUDY compared to the original BarFLEX collection (Douglas, et al. 2012). 70% of them are not annotated in the Saccharomyces Genome Database (SGD) and thus have not been experimentally characterized. (D) Assessing the impact of increased expression on colony growth in two non-stress and stress environment pairs: (1) SC-URA+galactose (“SC”) and SC-URA+galactose+NaCl (“NaCl”); (2) SC-URA+galactose+DMSO (“DMSO”) and SC-URA+galactose+tunicamycin (“tunicamycin”). DMSO is the vehicle of the stressor tunicamycin. (E) Growth scores in the SC environment in this study correlate well with measurements from published work (Vakirlis, et al. 2020). ORF: open reading frame.

To this aim, we deployed a new yeast overexpression collection containing nearly a thousand evolutionarily young *de novo* translated ORFs and systematically measured the overexpression phenotypes under stress and non-stress environments **(Figure 1B-D) (Methods)**. Follow-up experiments uncover that overexpression of *de novo* translated ORF YLR112W promotes growth in the presence of the drug rapamycin through large-scale transcriptomic rewiring. We subsequently named YLR112W *De novo Rapamycin response Dampener 1* (*DRD1*). The large-scale phenomic dataset provides insight into the potential of previously uncharacterized *de novo* translated ORFs to confer adaptive function after evolutionary changes and sheds light on the role of environmental context in determining the evolutionary fate of nascent proto-genes.

## Results

### Experimental simulation of *de novo* translated ORF evolution in stress environments

To assess the impact of increased *de novo* translated ORF expression on growth phenotypes, and whether or how such effects are modified by environmental conditions, we devised a yeast overexpression collection and deployed a high-throughput screening strategy to measure overexpression phenotype across different stress environments **(Figure 1B-D)**. As most *de novo* translated ORFs are neither annotated in the Saccharomyces Genome Database (SGD) (Engel, et al. 2025) nor included in any publicly available yeast collection (Carvunis, et al. 2012; Wacholder, et al. 2023; Rich, et al. 2024), we expanded one of the largest existing yeast overexpression collections, the BarFLEX collection (Douglas, et al. 2012) to cover hundreds of previously uncharacterized translated ORFs **(Figure 1B)**. This collection, which we named BUDY (<u>B</u>arFLEX with <u>U</u>ncharacterized *<u>D</u>e novo* ORFs in <u>Y</u>east), consists of ORFs with previously reported *de novo* origins (Carvunis, et al. 2012; Wacholder, et al. 2023; Rich, et al. 2024) and native translation (Wacholder, et al. 2023). The overexpression is driven by a galactose-inducible promoter on a Yeast Centromere (CEN) plasmid as in the original BarFLEX collection (Douglas, et al. 2012). To estimate cloning success for the BUDY collection, we verified the sequences in the plasmids for 115 arbitrarily selected strains, including 102 from the expanded part of the collection. As expected, most of the strains (104 out of 115) have the correct sequences **(Table S1)**. We therefore inferred a correct genotype rate of 90.4% for the BUDY collection. The strains with incorrect sequences were filtered out. In total, 5,083 translated ORFs were considered for experiments and analyses **(Figure 1B and Table S1)**. 893 out of the 5,083 translated ORFs are new in the BUDY collection. Among all 5,083 translated ORFs, 945 have evidence of *de novo* origin within the *Saccharomyces* genus (Carvunis, et al. 2012; Wacholder, et al. 2023; Rich, et al. 2024), whereas 4,138 are conserved, defined as being present outside the genus or showing a high level of reading frame conservation within the genus (Carvunis, et al. 2012; Wacholder, et al. 2023; Rich, et al. 2024). Out of the 945 *de novo* translated ORFs, 663 are not annotated in the SGD, making this study the first experimental work for these *de novo* translated ORFs. The BUDY collection is, to our knowledge, the largest for *de novo* ORF overexpression in any species **(Figure 1C)**.

In addition to the overexpression strains, the BUDY collection also contains an empty-plasmid reference strain for comparison. This enables the calculation of growth scores, defined here as the ratio of the colony size of overexpression strains to the colony size of the reference strain after spatial correction **(Figure 1D, Methods)**. We measured growth scores of individual translated ORF overexpression in four types of media based on synthetic complete (SC) media lacking uracil (-URA, for plasmid retention) and with galactose to induce overexpression without and with stressors **(Table S2)**: SC-URA (hereafter "SC"), SC- URA+DMSO ("DMSO"), SC-URA+NaCl ("NaCl"), and SC-URA+tunicamycin ("tunicamycin"). The first two are the respective non-stress controls of the last two environments. NaCl (osmotic stress) and tunicamycin (endoplasmic reticulum (ER) stress via unfolded protein response) were selected as they are commonly used in high-throughput genetic screens (Turco, et al. 2023). The SC and NaCl environments were repeated to assess reproducibility. The two experiments correlate well with each other in the growth scores **(Supplementary Figure 1A)**. In addition, the growth scores in the SC-URA environment correlate with previous work using colony-based phenotyping **(Figure 1E and Supplementary Figure 1B)** (Vakirlis, et al. 2020) and liquid-based pooled assays **(Supplementary Figure 1B)** (Douglas, et al. 2012). Together, these results establish that our colony-based growth scores are reproducible across experiments and concordant with an orthogonal phenotyping approach.

### Osmotic and ER stresses often positively modify the phenotypic impact of increased ***de novo* translated ORF expression**

To investigate whether, and how, environments affect the overexpression phenotypes of *de novo* translated ORFs, we first compared the growth scores among environments for each strain group **(Figure 2A)**. We found that growth scores of the *de novo* translated ORF group varied across environments, similar to the behavior of the conserved translated ORF group. As expected, such variation did not exist in the reference strain across environments. This suggests that the phenotypic impact of increased expression of *de novo* translated ORF as a group depends on environment.

**Figure 2.**
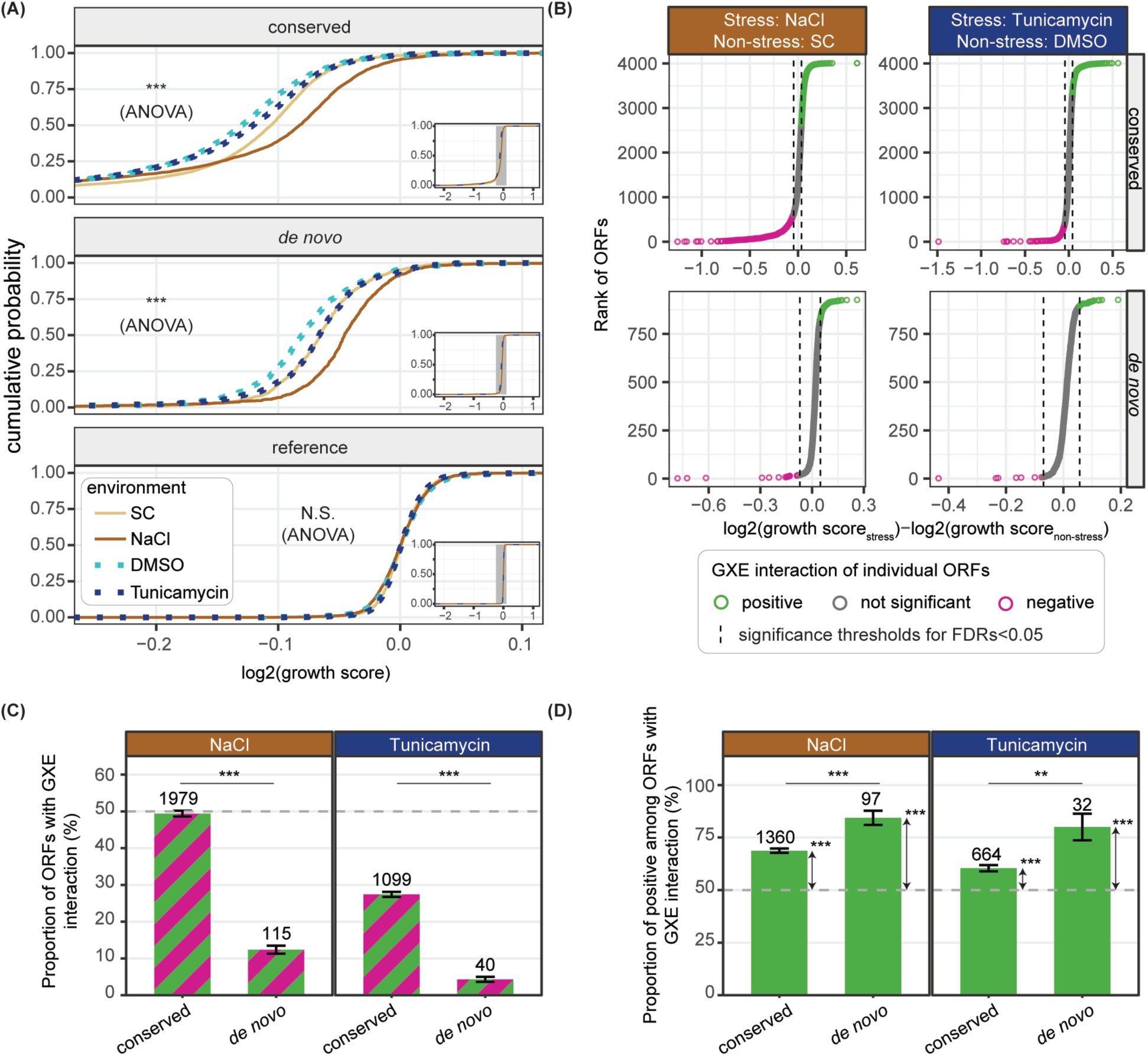
Phenotypes of increased *de novo* translated ORF expression depend on environmental contexts. (A) Cumulative distributions of growth scores of each strain type in four environments. The environment affects the overexpression phenotypes of both conserved and *de novo* translated ORF overexpression strains (ANOVA). As expected, growth scores of the reference strain do not vary among environments (ANOVA). The main panels are the zoomed-in views of the shaded range in the insets. (B) GXE interactions of ORFs based on their phenotypes in paired stress and non-stress environments (NaCl vs. SC; tunicamycin vs. DMSO). Translated ORFs with significantly higher growth scores under stress have positive GXE interactions (green); those with lower scores have negative interactions (magenta). GXE interactions were detected for each combination of evolutionary class and environment separately while maintaining the empirical FDRs under 5%. The dashed lines are the significance thresholds. (C) Proportions of conserved and *de novo* translated ORFs with significant GXE interactions with NaCl or tunicamycin, regardless of direction. GXE interactions are more common among conserved than *de novo* translated ORFs for both stressors (χ^2^ test). (D) Among translated ORFs with significant interactions, positive interactions are more prevalent in *de novo* than conserved translated ORFs for both stressors (χ^2^ test, visualized as the deviation from 50%, which is indicated by the dashed lines). Such bias towards positive interaction is stronger in the *de novo* group (χ^2^ test). In (C) and (D), numbers above the bars are translated ORF count. Error bars are the standard error of proportions. GXE interaction: gene-by-environment interaction. N.S.: not significant. **: p<0.01. ***: p<0.001. SC: SC-URA+galactose, DMSO: SC-URA+galactose+DMSO, NaCl: SC-URA+galactose+NaCl, tunicamycin: SC-URA+galactose+tunicamycin.

To further investigate the environmental dependency of overexpression phenotype for individual *de novo* translated ORFs, we identified translated ORFs whose overexpression phenotype was modified by either stressor, NaCl and tunicamycin, using gene-by-environment (GXE) analysis **(Methods)**. We considered a translated ORF-stressor pair to have a positive GXE interaction when a translated ORF had a higher growth score in a stress environment than in the corresponding non-stress control environment, meaning the stress improves the impact of increased expression, and vice versa for negative interactions. We detected GXE interactions in both directions for both conserved and *de novo* translated ORFs **(Figure 2B and Table S3)**. In total, 12.4% and 4.3% of the *de novo* translated ORFs tested showed significant GXE interactions with NaCl and tunicamycin, respectively. These proportions were lower compared to those of conserved translated ORFs **(Figure 2C)**. The results demonstrate that the phenotypic impact of increased expression is modified by environmental stresses for a non-negligible proportion of *de novo* translated ORFs.

The direction of GXE interactions is particularly informative for understanding how stress shapes *de novo* gene birth: a positive interaction suggests that stress makes increased translated ORF expression more tolerable or beneficial, whereas a negative one suggests the opposite. We found that positive interactions outnumbered negative ones for both NaCl and tunicamycin among *de novo* translated ORFs, and this directional bias was significantly stronger in *de novo* translated ORFs than in conserved ones **(Figure 2D)**. Thus, these stressors more likely improve than worsen the phenotypic impact of increased *de novo* translated ORF expression. This is consistent with Fisher’s geometric model, which predicts that organisms farther from their fitness optimum are more likely to tolerate or even benefit from mutations (Fisher 1930; Orr 2005).

### Stress severity positively modifies phenotypic impact of increased *de novo* translated ORF expression

To explore whether environmental stresses generally positively modify the phenotypic impact of increased *de novo* translated ORF expression, we expanded the environmental panel to 22 different conditions and measured the growth scores of a subset (N=557) of the 945 *de novo* translated ORFs in the BUDY collection. This subset is mostly comprised of *de novo* ORFs **(Figure 1)** supported by multiple studies **(Table S4)** (Carvunis, et al. 2012; Wacholder, et al. 2023; Rich, et al. 2024). These 22 environments span from high (39°C) and low (17°C) temperatures, drugs that target different pathways (such as DNA synthesis, membrane synthesis, and Target of Rapamycin (TOR) pathways), various pH to nutrient limitations in addition to osmotic and ER stresses **(Figure 3 and Table S5)**.

**Figure 3.**
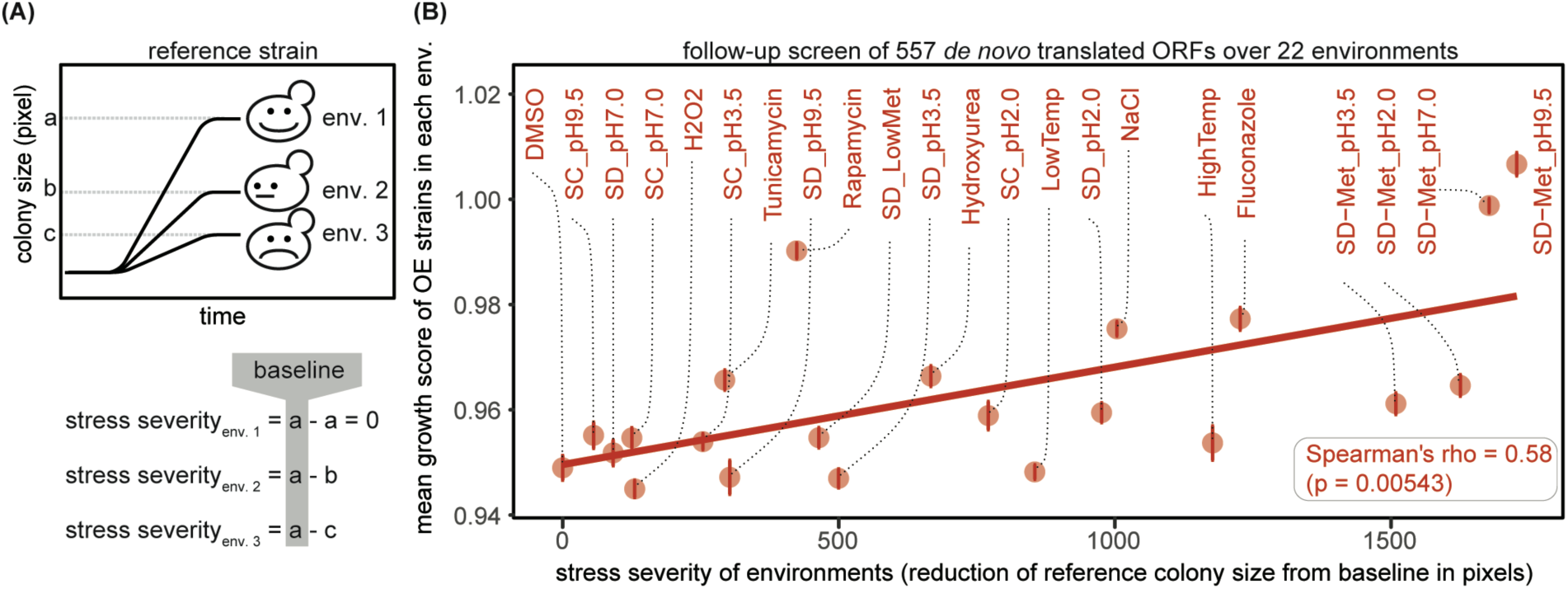
Stress severity positively correlates with mean *de novo* translated ORF overexpression growth scores across 22 environments. (A) Conceptual illustration of stress severity measurement. Stress severity was calculated by subtracting the maximum reference colony size of each environment from that of the least stressful environment; higher values indicate greater stress. (B) Scatter plot of stress severity against mean growth score across environments in the follow-up screen with 557 *de novo* translated ORFs over 22 environments. Error bars indicate the standard error of the mean. Only translated ORFs tested in at least 10 environments are included. All media are based on SC-URA+galactose except for those with the prefix “SD”, which are based on SD-URA+galactose.

To quantify stress severity across environments, we leveraged the maximal colony sizes of the reference strain as a proxy for stress severity, as growth curves plateau at smaller colony sizes under greater stress **(Figure 3A)**. The stress severity exhibited a significant positive correlation with the mean growth score of increased *de novo* translated ORF expression (Spearman’s rho = 0.580, p = 0.005) **(Figure 3B)**. As expected, the correlation was not statistically significant in the reference strain (p=0.086) and the correlation of *de novo* translated ORFs differed from that of the reference strain (likelihood-ratio test, p < 1.678x10^-15^). These results provide further evidence that the phenotypic impact of increased *de novo* translated ORF expression is positively modified by environmental stress, such that severe stresses appear more permissive for increased *de novo* translated ORF expression.

### Beneficial *de novo* translated ORFs in specific stresses

Having established that environmental stresses positively modify the phenotypic impact of increased *de novo* translated ORF expression as a group, a key follow-up question is whether individual *de novo* translated ORFs can confer beneficial phenotypes upon increased expression, and if so, in which environments. We compared the growth scores of each translated ORF to the reference strain in each environment and discovered beneficial or deleterious phenotypes after stringent empirical FDR control for either direction in each environment **(Table S5 and S6)**. We aimed to control the empirical FDRs under 0.05 **(Methods)**. The achieved empirical FDRs were substantially more stringent than the 0.05 target in all environments for beneficial phenotypes and in most environments (15/22) for deleterious phenotypes, approaching 0 in many cases **(Table S6)**. The proportion of translated ORFs with significant phenotypes varied across environments **(Figure 4A)**. In total, 485 out of 557 *de novo* translated ORFs showed phenotypes in either or both directions across environments. Among them, 30 *de novo* translated ORFs showed beneficial phenotypes, and 480 showed deleterious phenotypes in at least one environment. These numbers constituted around 5.4% and 86.2% of the translated ORFs tested in this study. The 30 beneficial translated ORFs improved growth scores by 3.7 to 38.8% (mean=17.5%; median=16%). These results show that, despite being outnumbered by deleterious translated ORFs, a substantial proportion of *de novo* translated ORFs can produce a beneficial phenotype upon increased expression in an environment-dependent manner, confirming that *de novo* translated ORFs have the potential to mediate adaptive evolution in specific environments (Vakirlis, et al. 2020).

**Figure 4.**
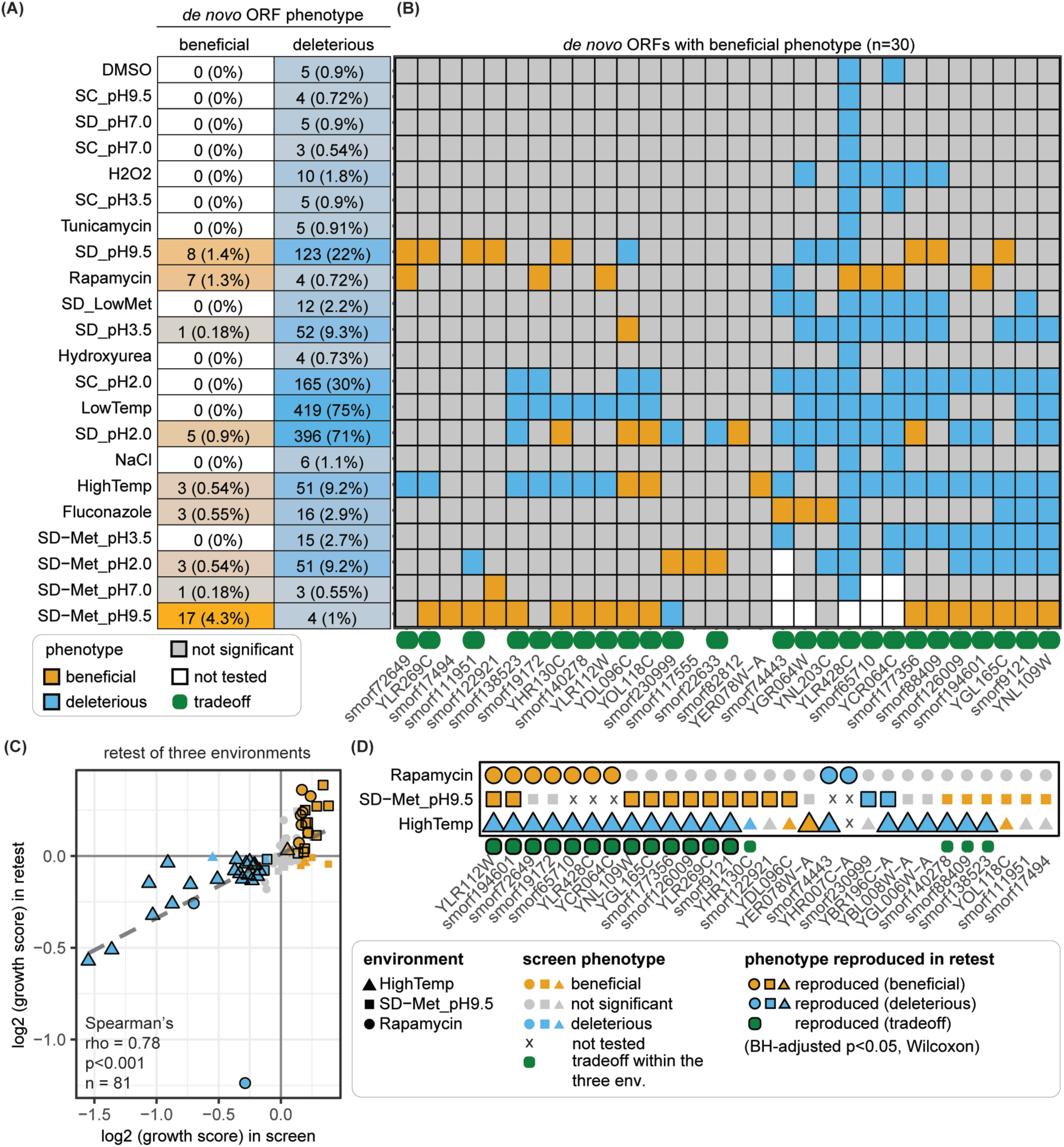
Phenotypes of increased expression of individual *de novo* translated ORFs reveal environmental dependency and phenotypic tradeoffs across 22 environments. (A) Numbers and proportions of *de novo* translated ORFs showing beneficial or deleterious phenotypes in each environment upon overexpression. Parentheses indicate the proportion of translated ORFs with detected phenotypes among all translated ORFs tested in that environment. The proportions of beneficial and deleterious phenotypes vary across environments. (B) Phenotypic tradeoffs are prevalent among the 30 beneficial translated ORFs (columns). Most (25/30; 83.3%) also showed deleterious phenotypes in at least one environment. Green boxes indicate translated ORFs with phenotypic tradeoffs. 29 *de novo* translated ORFs with beneficial and/or deleterious phenotypes were selected for a retest over three environments. (C) and (D) show results of the retest of 29 beneficial or deleterious translated ORFs. In the retest, the 29 strains were retaken from the glycerol stock and retested over three environments. (C) Phenotypes of 27 translated ORFs in the retest compared to the main screen. Black outlines indicate phenotypes reproduced in the retest. Note that 29 translated ORFs were retested, two of which were not tested in the screen for a few environments. (D) Phenotypes of 29 translated ORFs in three environments in retest. All media are based on SC-URA+galactose except for those with the prefix “SD”, which are based on SD-URA+galactose.

### Phenotypic tradeoff is prevalent among beneficial *de novo* translated ORFs

An extreme scenario of environmental dependency of phenotypic impact caused by increased expression is phenotypic tradeoff, namely, causing beneficial phenotypes in some environments while causing deleterious phenotypes in other environments. Phenotypic tradeoff is of particular interest because the direction of natural selection on increased expression of translated ORFs with tradeoff is presumably dependent on the environment. Therefore, we investigated whether the 30 *de novo* translated ORFs had phenotypic tradeoffs by examining their phenotype across the 22 environments **(Figure 4B)**. Most of the beneficial *de novo* translated ORFs (25 out of 30; 83.3%) also showed deleterious phenotypes in other environments. The prevalent phenotypic tradeoff among these *de novo* translated ORFs underscores the important role of environments in determining the overexpression phenotypes of *de novo* translated ORFs.

To assess the reproducibility of *de novo* translated ORF overexpression phenotypes and their associated tradeoffs, we performed an independent retest in three environments: rapamycin, SD-MET pH9.5, and high temperature. We chose the first two environments for their high numbers of beneficial translated ORFs in the screen described in the previous paragraph (hereafter “the screen”). The high temperature environment was chosen to maximize the number of translated ORFs with tradeoff in the retest. We retested 29 translated ORFs with beneficial and/or deleterious phenotypes in the three environments, allowing the measurement of growth scores for a total of 87 environment-translated ORF pairs, 52 of which showed beneficial or deleterious phenotypes in the screen **(Table S7)**. We picked fresh cells from the glycerol stocks of the BUDY collection, and measured growth scores again with the same colony-based assay. We found that the growth scores measured in this independent retest correlated well with the original measurement from the screen **(Figure 4C)** (Spearman’s rho=0.78, p<0.001). Notably, 43 out of 52 environment-translated ORF pairs exhibited consistent beneficial or deleterious phenotypes between the retest and the screen **(Figure 4C-D and Supplementary Figure 2-4)** (Benjamini-Hochberg (BH)-adjusted p values< 0.05, Wilcoxon Rank Sum test). Among the 17 retested translated ORFs that showed phenotypic tradeoff in the screen, 13 showed phenotypic tradeoffs in the retest **(Figure 4C-D and Supplementary Figure 2-4)**. Therefore, our independent retest experiment confirms that the phenotypes mediated by increased *de novo* translated ORF expression that we observed in our screens, and their associated tradeoffs, are robust.

### Using YLR112W to gain mechanistic insight into its beneficial overexpression in rapamycin stress

To explore the molecular mechanisms underlying how the phenotypic effects of increased *de novo* translated ORF expression depend on environmental conditions, we focused on YLR112W overexpressed under rapamycin stress for a case study. This translated ORF- stressor pair was selected for its reproducible increased growth score phenotype **(Figure 4D)**, and the high-confidence *de novo* evolutionary origin of the translated ORF as revealed by a previous case study (Lu, et al. 2017). We named this ORF *De novo Rapamycin response Dampener 1* (*DRD1*).

Rapamycin inhibits growth by inducing a growth-repressing transcriptome program even when the environment is otherwise favorable for growth (De Virgilio and Loewith 2006; Rohde, et al. 2008; González and Hall 2017; Foltman and Sanchez-Diaz 2023). Therefore, we aimed to use RNA-seq to investigate the impact of Drd1 overexpression on the transcriptomic response to rapamycin. We designed and constructed new strains **(Figure 5A)** for follow-up experiments with several key technical changes to facilitate the RNA-seq experiment as well as to assess how robust the beneficial phenotype is to technical choices. These technical changes include: (1) a C-terminal mNeonGreen (mNG) tag to allow confirmation of overexpression at the protein level; (2) Z3EV promoter-based β-estradiol-inducible overexpression system (McIsaac, et al. 2013), instead of galactose-inducible system, allowing the more common carbon source glucose instead of galactose as the carbon source; (3) genomic integration of the overexpression construct instead of plasmid-based overexpression; and (4) liquid growth assay instead of colony-based assay.

**Figure 5.**
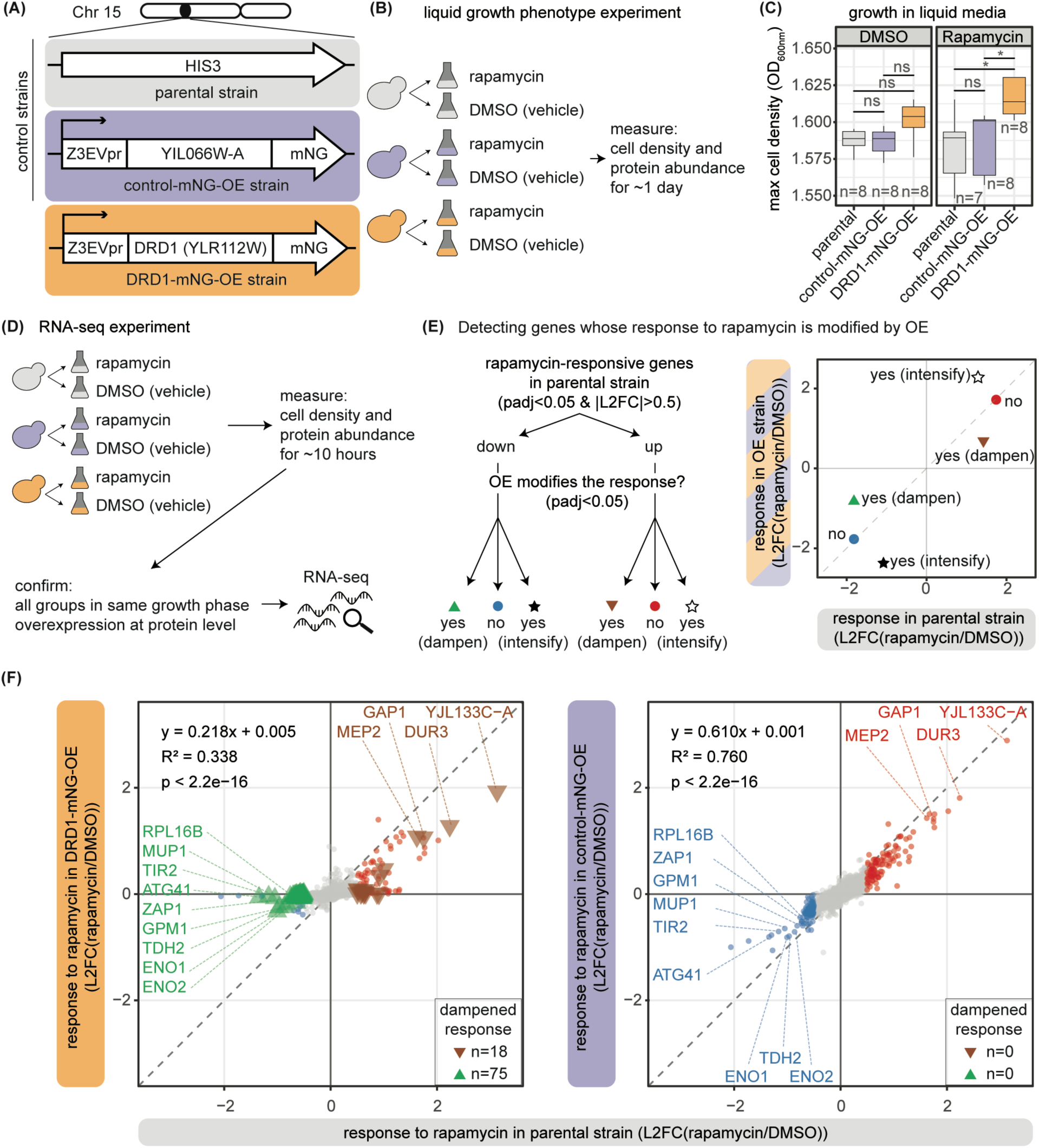
*DRD1* (YLR112W) overexpression dampens the growth-inhibiting transcriptomic program induced by rapamycin. (A) Strain design for the follow-up investigation of *DRD1*. Key differences from the BUDY cloning system include genomic integration instead of plasmid-based expression, use of the Z3EV promoter, and addition of a C-terminal mNeonGreen tag to enable protein abundance tracking **(Methods)**. YIL066W-A was selected arbitrarily from *de novo* translated ORFs without significant phenotype in the colony-based screen and was used as a neutral control ORF for comparison with *DRD1*. (B) Workflow of phenotyping experiment in liquid media with the new strains. (C) Maximum cell densities in a liquid growth assay with media with the stressor rapamycin or its vehicle DMSO. *: BH-adjusted p < 0.05, Wilcoxon Rank Sum test. (D) Workflow of the RNA-seq experiment. (E) Analytical pipeline and conceptual illustration of the detection of genes whose response to rapamycin is modified by overexpression. (F) Drd1 overexpression dampens the transcriptomic response to rapamycin. The effect of the neutral ORF overexpression is shown for comparison. No gene with an intensified response was detected in either comparison. Pearson correlation coefficients and linear model fits are shown. Insets indicate the number of genes with a significant impact on the response. Gene labels: the intersection of the 15 most up- or down-regulated genes in response to rapamycin in the parental strain and the genes with response dampened by Drd1 overexpression. Media are based on SC+glucose+β-estradiol, with the addition of either rapamycin or its vehicle DMSO. mNG: mNeonGreen. RPM: rapamycin. L2FC: log2 fold change. OE: overexpression.

In total, three test and control strains were constructed. They include: (1) a strain overexpressing C-terminally mNG-tagged Drd1 (hereafter “DRD1-mNG-OE strain”); (2) a strain overexpressing mNG-tagged Yil066w-a (hereafter “control-mNG-OE strain”), which is a neutral control translated ORF without overexpression phenotype in rapamycin stress in the screen; and (3) their parental strain without the genomic integration of the overexpression construct (hereafter “parental strain”) **(Figure 5A)**.

### Beneficial Ylr112w/Drd1 overexpression in rapamycin stress persists with orthogonal assay and a different expression system

To confirm that the overexpression occurs at protein level and that the DRD1-mNG-OE strain has a rapamycin-specific beneficial phenotype, we performed a liquid growth assay with SC+glucose+β-estradiol (for overexpression induction) media containing either rapamycin or its vehicle DMSO **(Figure 5B)**. As expected, both DRD1-mNG-OE and control- mNG-OE strains showed robust mNeonGreen signals **(Supplementary Figure 5A)**, confirming the overexpression at the protein level. We compared the maximum growth rates and maximum cell densities of the strains **(Figure 5C and Supplementary Figure 5B-C)**. Although no significant difference in maximum growth rates was detected, the DRD1-mNG- OE did achieve a higher maximum cell density than the two controls in rapamycin stress **(Figure 5C)**, showing the robustness of the beneficial Ylr112w overexpression phenotype in rapamycin stress despite changes in experimental setup.

### Ylr112w/Drd1 overexpression dampens the growth-repressing transcriptomic response to rapamycin

Having established that Drd1 has a robust beneficial overexpression phenotype in rapamycin stress, we performed RNA-seq experiments to investigate the impact of Drd1 overexpression on the growth-inhibiting transcriptomic response to rapamycin using the DRD1-mNG-OE strain and its controls, the parental and the control-mNG-OE strains **(Figure 5A, 5D-E, Supplementary Figure 6, and Table S8)**. We collected cells at a time point when all three strains were near the late exponential phase to ensure comparability, and that DRD1-mNG-OE and control-mNG-OE strains showed fluorescent signals **(Supplementary Figure 6) (Methods)**. As expected, the RNA-seq data further confirmed the overexpression at the RNA level **(Supplementary Figure 7A)** and samples of each of the genotype-environment groups were clustered together in terms of their transcriptomic profiles **(Supplementary Figure 7B)**, confirming that overexpression and/or the drug did impact the transcriptome.

Rapamycin is a natural growth inhibitor (Heitman, et al. 1991; Powers and Kellogg 2022). It forms a complex with FKBP12, which subsequently binds and inhibits the target of rapamycin (TOR) pathway, a central regulatory hub of growth, resulting in growth-repressing, “starvation-like” transcriptomic responses, such as down-regulation of genes involved in translation, ribosome biogenesis, and up-regulation of permeases for nutrient scavenging, even if the environment is otherwise favorable for growth (De Virgilio and Loewith 2006; Rohde, et al. 2008; González and Hall 2017; Foltman and Sanchez-Diaz 2023).

Comparison of the transcriptome of the parental strain in rapamycin stress to that in the medium with DMSO (hereafter “rapamycin” and “DMSO” respectively) provides a baseline of transcriptomic response to rapamycin without overexpression **(Figure 5D and Supplementary Figure 8A and Table SG)**. We detected differentially expressed genes (DEGs, genes with adjusted p < 0.05 and |log_2_ fold change| > 0.5) in response to rapamycin in both directions in the parental strain, including 109 and 130 genes up- and down- regulated DEGs, respectively. As expected, genes related to transport and translation were among the most up- and down-regulated rapamycin-responsive genes of the parental strain, respectively **(Supplementary Figure 8A)**. Consistently, biological processes known to respond to rapamycin were enriched in the DEGs in either direction, including ribosomal protein biosynthesis, translation initiation, and rRNA processing among the down-regulated DEGs, and nitrogen catabolism, stress response, and autophagy among the up-regulated DEGs (Hardwick, et al. 1999; Shamji, et al. 2000) **(Supplementary Figure 8B)**. Gene targets of transcription factors (TFs) downstream of the TOR pathway were also enriched in the DEGs **(Supplementary Figure 8B)**. These TFs include *RGT1*, a regulator of glucose transporter, among the up-regulated DEGs, and *HMO1*, *RAP1,* and *IFH1*, key regulators of ribosome biogenesis, among the down-regulated DEGs (De Virgilio and Loewith 2006; Rohde, et al. 2008; González and Hall 2017; Foltman and Sanchez-Diaz 2023). Altogether, the transcriptomic response to rapamycin in the parental strain recapitulates prior knowledge of the growth-repressing transcriptomic rewiring induced by rapamycin.

To examine the impact of Drd1 overexpression on transcriptomic response to rapamycin, we compared the transcriptomic responses to rapamycin, measured as the log2 fold change in response to rapamycin (hereafter “L2FC(RPM/DMSO)”), of either DRD1-mNG-OE or control-mNG-OE strains to that of the parental strain **(Figure 5D-F and Table S10)**. In theory, if a translated ORF overexpression does not affect the response to rapamycin of a gene, the L2FC(RPM/DMSO) of this gene in the overexpression strain and the parental strain should be similar **(Figure 5E)**. In contrast, if the overexpression dampens or intensifies the rapamycin response of a gene, the L2FC(RPM/DMSO) should differ between the overexpression strain and the parental **(Figure 5E**). As expected, the L2FC(RPM/DMSO) of the control-mNG-OE strain showed a high correlation with that of the parental strain (R^2^=0.76, Pearson correlation, p<2.2x10^-16^, slope=0.610) **(Figure 5F, right)**. In contrast, we observed a poor correlation of L2FC(RPM/DMSO) between the DRD1-mNG-OE strain and the parental control strain (R^2^= 0.338, Pearson correlation, p<2.2x10^-16^, slope=0.218) **(Figure 5F, left)**. Furthermore, the slope in the comparison of the DRD1-mNG-OE strain to the parental strain is significantly reduced from that of the control-mNG-OE strain to the parental strain (p < 2x10^-16^). These results demonstrate that YLR112W overexpression dampens the transcriptomic response to the growth inhibitor rapamycin.

To understand the dampening effect of Drd1 overexpression in greater detail, we identified the specific genes whose response to rapamycin is modified by Ylr112w overexpression using GXE interaction analysis **(Figure 5F and Table S10) (Methods)**. Of the 109 and 130 genes up- and down-regulated by rapamycin in the parental strain, 18 (16.5%) and 75 (57.7%), respectively, showed dampened rapamycin response in the DRD1-mNG-OE strain. The dampening effect was proportionally more common among the genes down-regulated by rapamycin in the parental strain (p=1.888x10^-10^, χ^2^ test). As expected, overexpression of the neutral ORF, Yil066w-a, did not significantly modify the response of any genes **(Figure 5F)**. In addition, genes with dampened up- or down-regulation in response to rapamycin span a wide range of biological processes and are downstream to a wide range of TFs regulated by the TOR pathway **(Figure 5F and Supplementary Figure 8C)**, suggesting that Ylr112w overexpression confers a beneficial phenotype in rapamycin stress through extensive dampening of growth-repressing transcriptomic responses.

## Discussion

In this study, we demonstrate that the phenotypic impacts of increasing expression of *de novo* translated ORFs vary substantially with environmental conditions. Our gene-by-environment interaction analyses revealed that notable proportions of *de novo* translated ORFs exhibit environment-dependent overexpression phenotypes and that stressors are more likely to improve than to worsen the phenotypic impact of increased expression of *de novo* translated ORFs. We observed a strong positive correlation between environmental stress severity and the mean phenotypic impacts of increased *de novo* translated ORF expression. Notably, we identified 30 *de novo* translated ORFs that confer beneficial phenotypes in specific environments, with most (25/30) exhibiting deleterious phenotypes in other environments. Focusing on Ylr112w, renamed Drd1, for a case study, we uncovered that when its expression is increased, this *de novo* ORF mediates a large-scale transcriptomic rewiring that confers resistance to the growth inhibitor rapamycin. Together, these results demonstrate that environmental stresses profoundly modulate the direction and magnitude of phenotypic impact of increased *de novo* translated ORF expression and reveal transcriptomic rewiring as one underlying molecular mechanism of environment-specific beneficial *de novo* translated ORF overexpression.

### Implication in dark proteome research and beyond

Recent omics studies have revealed numerous unannotated ORFs with native translation (Ingolia, et al. 2009; Carvunis, et al. 2012; Ruiz-Orera and Albà 2019; Mudge, et al. 2022; Wacholder, et al. 2023; Chothani, et al. 2026; Deutsch, et al. 2026). Most of these sequences have not been functionally characterized. These sequences, often referred to as the “dark translatome” or “dark proteome” (Wright, et al. 2022; Casola, et al. 2025), are mostly comprised of translated ORFs of recent evolutionary origins (Sandmann, et al. 2023; Wacholder, et al. 2023). In total, 663 *de novo* translated ORFs overexpressed in this current study are unannotated **(Figure 1C)**, making this study of *de novo* ORFs directly relevant to dark proteome or translatome research.

It has been proposed that *de novo* translated ORFs are a source of new genes and lineage-specific traits (Van Oss and Carvunis 2019; Ardern 2023; Ardern and uz-Zaman 2023; Zhao, et al. 2024; Barrera-Redondo, et al. 2025; Chou, et al. 2026), as they acquire gene-like properties, such as increased expression levels, during evolution (Carvunis, et al. 2012). This current study fits into this bigger picture by experimentally demonstrating beneficial phenotypic consequences of the increased expression of individual *de novo* translated ORFs, many of which are unannotated. This supports the idea that some of these sequences have adaptive potential, i.e., the ability to confer adaptive function, once they acquire gene-like properties during evolution. The fact that some sequences of the dark proteome have adaptive potential emphasizes the need to expand the scope of research on the dark proteome: besides asking what biological function the dark proteome confers at current evolutionary time point, it is also important to investigate its potential to give rise to new genes and promote organismal adaptation in the future.

In addition, there are a handful of large-scale studies that performed CRISPR knockout screens on hundreds or thousands of ORFs that are not annotated as protein-coding in databases using different human cell lines (Chen, et al. 2020; Prensner, et al. 2021; Wacholder, et al. 2023; Zheng, et al. 2023; Hofman, et al. 2024; Pai, et al. 2025; Schlesinger, et al. 2025; Valdivia-Francia, et al. 2025). It is possible that many *de novo* translated ORFs were tested in these experiments. We envision that reanalysis of these genetic screen datasets of unannotated translated ORFs after filtering for those with *de novo* origins may be complementary to our study in terms of species and direction of genetic manipulation (knockout vs overexpression). A successful example of such reanalysis was done by Vakirlis et al. They found that many unannotated translated ORFs with deletion phenotypes in another prior CRISPR-based genetic screen study (Chen, et al. 2020) have *de novo* evolutionary origins (Vakirlis, et al. 2020), underscoring the utility of these large-scale genetic screen studies of unannotated translated ORFs in the context of *de novo* gene birth research.

More generally, genetic variations in noncoding genomic regions, or the "dark genome," in a population are associated with adaptive traits, as often revealed by quantitative trait loci mapping (QTL mapping) (She and Jarosz 2018; Nguyen Ba, et al. 2022). These genetic variations in noncoding regions may sometimes be explained by their effects on the expression or regulation of protein-coding genes. Alternatively, some of these genetic variations may reside in or near regions that encode the dark proteome, conferring phenotypic effects through these previously overlooked coding sequences. We envision that the unannotated ORFs with overexpression phenotypes found in our current study can serve as a starting point to revisit trait-associated genetic variations in “noncoding” regions, namely, whether they affect phenotype through increasing the expression of *de novo* and/or unannotated translated ORFs.

### How environmental stress affects the progression of de novo gene birth

Environmental stress has long been suspected to play a critical role in *de novo* gene birth (Carvunis, et al. 2012; Schlötterer 2015). Across diverse species, *de novo* translated ORFs are frequently upregulated in response to environmental stresses (Schlötterer 2015), raising the possibility that some of these sequences already contribute, or have the potential to evolve into *de novo* genes that contribute, to stress adaptation. Sporadic case studies further support this view, such as the *de novo* antifreeze glycoprotein in Arctic cod, whose expression enables survival in freezing environments (Baalsrud, et al. 2018; Zhuang, et al. 2019) and the *de novo* gene that increases drought tolerance of seeds in *Arabidopsis* (Jin, et al. 2025). Yet it is unclear whether or how environmental factors steer the progression of *de novo* gene birth. Our systematic examination of the phenotypic impact of increased *de novo* translated ORF expression sheds light on this question. We show that osmotic and ER stresses overwhelmingly bias GXE interactions in the positive direction, indicating that these stressors are more likely to improve than to worsen the phenotypic impact of increased *de novo* translated ORF expression. Extending this analysis to 22 diverse environments, we find a strong positive correlation between stress severity and the mean phenotypic impact of increased expression. Together, these results raise the possibility that environments of higher stress severity may be more permissive for the evolutionary exploration of higher expression levels of these novel sequences. This observation is conceptually aligned with classical evolutionary theory, namely Fisher’s geometric model, which predicts that organisms farther from their fitness optimum, such as under stress, are more likely to tolerate or benefit from mutations (Fisher 1930; Orr 2005). By analogy, increased expression of a *de novo* translated ORF that is neutral or deleterious under optimal growth may become beneficial or less harmful in challenging environments, thereby increasing the likelihood that elevated expression levels are retained by selection. Together, our results suggest a positive role of environmental stress in facilitating the progression of *de novo* gene birth.

### Phenotypic tradeoffs across environments as a potential contributor to fast evolutionary turnover of de novo sequences

Previous studies suggest that incipient emergence and increased expression of *de novo* translated ORFs can confer beneficial phenotypes that favor their selection (Neme, et al. 2017; Vakirlis, et al. 2020; Castro and Tautz 2021; Bhave and Tautz 2022). According to this notion, one would expect that the genome of a species can become inflated by *de novo* genes over time. However, previous comparative genomic studies of different species have found that most *de novo* translated ORFs have recent evolutionary origins within genus, species, or populations, suggesting rapid evolutionary turnover of *de novo* translated ORFs (Tautz and Domazet-Lošo 2011; Wissler, et al. 2013; Neme and Tautz 2014; Palmieri, et al. 2014; Schlötterer 2015; Neme and Tautz 2016; Schmitz, et al. 2018; Durand, et al. 2019; Van Oss and Carvunis 2019; Heames, et al. 2020; Zheng and Zhao 2022; Grandchamp, et al. 2023; Roginski, et al. 2024; Zhao, et al. 2024). So, why do most *de novo* translated ORFs not persist over longer evolutionary times? This study provides a potential explanation. The widespread phenotypic tradeoff, i.e., beneficial in some environments and deleterious in others, of *de novo* translated ORFs, suggests that environmental changes may likely change the direction of selection. In a more general context, recent studies based on empirical observation and experimental evolution have found that new mutations conferring short-term beneficial impact do not guarantee their long-term fixation due to widespread phenotypic tradeoff of these mutations across environments (Khristich, et al. 2025; Song, et al. 2025). It is possible that *de novo* translated ORF evolution also conforms to this general trend, mediating short-term adaptation to stress while not persisting long-term due to phenotypic tradeoff.

### Transcriptomic reprogramming as a mechanism of environment-specific beneficial de novo translated ORF overexpression

We found that transcriptomic rewiring underlies the mechanism of the beneficial Drd1/Ylr112w-overexpression in rapamycin stress. Rapamycin is a naturally occurring growth inhibitor that inhibits the TOR pathway (Heitman, et al. 1991; Powers and Kellogg 2022). Its complex with FKBP12 inhibits cell growth by inhibiting the Target of Rapamycin Complex 1 (TORC1) and subsequently activates and deactivates different downstream TF targets of TORC1, leading cells to adopt a growth-inhibiting transcriptomic program that resembles starvation response (De Virgilio and Loewith 2006; Rohde, et al. 2008; González and Hall 2017; Foltman and Sanchez-Diaz 2023). More specifically, TORC1 phosphorylates two major downstream kinase effectors, Sch9 and Tap42. When TORC1 is active in the absence of rapamycin, phosphorylated Sch9 activates downstream TFs to maintain the transcription of genes related to translation and ribosome biogenesis; meanwhile, phosphorylated Tap42 represses PP2A phosphatase (De Virgilio and Loewith 2006; Rohde, et al. 2008; González and Hall 2017; Foltman and Sanchez-Diaz 2023). In the presence of rapamycin, inactive TORC1 results in the deactivation of Sch9 and Tap42. Deactivation of Sch9 results in down-regulation of its downstream genes, and the deactivation of Tap42 derepresses PP2A, which activates the TFs and up-regulates genes involved in general stress response and amino acid transport (De Virgilio and Loewith 2006; Rohde, et al. 2008; González and Hall 2017; Foltman and Sanchez-Diaz 2023). As we observed that the Drd1 overexpression dampened both down- and up-regulation that are typically downstream of Sch9 and Tap42, we suspect that the impact of Drd1 overexpression may stem from exerting effects on TORC1 directly rather than through Sch9 or Tap42 signaling. Possible mechanisms, such as preventing drug-target engagement through drug efflux or inhibition of rapamycin-FKBP12 complex formation or action, will require experimental investigation in future studies. We envision that the generation of interactome data of Drd1 or phosphoproteomic data to explore the phosphorylation state of the TOR pathway proteins in the future can describe the mechanism in greater detail.

### Opportunities for future technical improvement

Several limitations exist in our experimental system, representing opportunities to capture biology unexplored in this current study. First, we used a galactose-inducible promoter for *de novo* translated ORF overexpression in the BUDY collection (Douglas, et al. 2012; Gligorovski, et al. 2023). Previous studies show that the relationship between fitness and expression levels of protein-coding genes is not always linear (Perfeito, et al. 2011; Rest, et al. 2013; Keren, et al. 2016; Siddiq, et al. 2024). It is possible that some *de novo* translated ORFs with no phenotypic impact in our expression system can confer phenotype at lower overexpression levels. Second, the high-throughput phenotyping experiments in this study use colony sizes as the proxy for growth. The use of colony size-derived metrics as a proxy for growth or fitness is a well-established methodology that has been widely used for two decades (Collins, et al. 2006; Tong and Boone 2006; Baryshnikova, et al. 2010; Dittmar, et al. 2010; Lawless, et al. 2010; Levin-Reisman, et al. 2010; Wagih, et al. 2013; Young and Loewen 2013; Bean, et al. 2014; Levin-Reisman, et al. 2014; Wagih and Parts 2014; Bischof, et al. 2016; Zackrisson, et al. 2016; Herricks, et al. 2017). The growth metrics derived from colony sizes have been shown to have reproducible and moderate, but not perfect, correlations with growth metrics derived from orthogonal assays, such as the liquid-based pooled growth assay (Blomberg 2011; Douglas, et al. 2012; Zackrisson, et al. 2016). This indicates that growth measurements based on solid and liquid media are indicative of each other and complementary. In our follow-up study of Drd1/YLR112W, we demonstrate that its beneficial overexpression in rapamycin stress qualitatively persisted after changes to a different expression system and to the liquid assay. As the natural habitat of yeasts is diverse (Liti 2015) and that evolution of expression level is likely a spectrum rather than binary (overexpression vs no overexpression), we envision that adding orthogonal liquid-based assays and using a tunable expression system in the future will be helpful to capture a comprehensive picture of the biology of *de novo* translated ORFs.

### Large-scale genetic screens in de novo gene birth research: comparison to prior work and opportunities

High-throughput experimentation represents a powerful approach to understanding *de novo* gene birth. Expressing random sequence libraries has been used to experimentally simulate the earliest stage of *de novo* gene birth – incipient expression of sequences with no prior exposure to natural selection in the form of proteins (Neme, et al. 2017; Knopp, et al. 2019; Castro and Tautz 2021; Knopp, et al. 2021; Bhave and Tautz 2022; Babina, et al. 2023; Frumkin and Laub 2023; Frumkin, et al. 2025). This approach, mostly based on *Escherichia coli*, has led to the discovery of various phenotypic impacts of random sequence expression, including growth improvement, rescuing auxotrophy, and resistance to antimicrobial drugs, bacterial toxins, and phages (Neme, et al. 2017; Knopp, et al. 2019; Castro and Tautz 2021; Knopp, et al. 2021; Bhave and Tautz 2022; Babina, et al. 2023; Frumkin and Laub 2023; Frumkin, et al. 2025), demonstrating that functional proteins can be born from scratch. The proportions of beneficial sequences in a random sequence pool vary greatly depending on studies and type of experimental strategy. Those studies that profile the phenotypes of individual mutants report beneficial rates ranging from 16% to 25% (Neme, et al. 2017; Castro and Tautz 2021; Bhave and Tautz 2022) whereas those studies that subject the random sequence pools to harsh selection condition to select for few sequences whose expression rescues growth report lower beneficial sequence rates ranging from 10^-8^ to 5x10^-5^ (Knopp, et al. 2019; Knopp, et al. 2021; Frumkin and Laub 2023; Frumkin, et al. 2025).

Direct testing of real *de novo* sequences, i.e., those encoded by genomes, is limited. A previous, proof-of-concept study used a small set of *de novo* ORFs and environments (Vakirlis, et al. 2020). In this previous study, all overexpressed ORFs were annotated in SGD and thus biased towards higher evolutionary conservation than the total pool of *de novo* translated ORFs in yeasts (Carvunis, et al. 2012; Wacholder, et al. 2023; Rich, et al. 2024). To our knowledge, the present study is by far the largest phenotypic study of increased *de novo* translated ORF expression in any species, in terms of both the number of sequences and the number of environments investigated. This current study discovers *de novo* translated ORFs with beneficial overexpression. It not only qualitatively confirms the finding from the previous proof-of-concept study but also demonstrates that beneficial overexpression extends to unannotated translated ORFs and that environmental context can change the magnitude and direction of overexpression phenotypes.

Around 5.4% of all *de novo* translated ORFs tested were beneficial in at least one environment in this current study, slightly lower than the previous estimate based on *de novo* sequences in yeast (10%) (Vakirlis, et al. 2020) and the estimates from studies that profiled the phenotypes of random sequence expression in *E. coli* (16-25%) (Neme, et al. 2017; Castro and Tautz 2021; Bhave and Tautz 2022). The difference might be explained by the more conservative statistical framework in this current study **(Methods)**, choice of species and assays, or types of sequences (*de novo* ORFs or yeast vs random sequence). The difference between this current study and the previous proof-of-concept yeast *de novo* sequence study (Vakirlis, et al. 2020) may also be due to the increased sequence set that covers translated ORFs beyond SGD annotation. Nonetheless, our overexpression collection still covers only a fraction of the full repertoire of recently evolved ORFs in *S. cerevisiae* (Carvunis, et al. 2012; Wacholder, et al. 2023; Rich, et al. 2024). A larger collection can help refine this estimate in the future. Overall, the scale of this study potentiates novel insights into environmental dependency, such as the discovery of widespread phenotypic tradeoffs among beneficial *de novo* translated ORFs. We envision that the phenotypic data generated in this study will be a valuable resource to the community for *de novo* gene birth research.

## Materials and methods

### Selection and classification of ORFs for the overexpression screens

ORFs were selected and classified for the overexpression screens based on their translation and evolution status. We first used a published comprehensive list of translated ORFs in yeast (Wacholder, et al. 2023) to select for ORFs with native translation. This list was made by large-scale integration of ribo-seq data and is by far the most comprehensive list of translated ORFs of yeast (Wacholder, et al. 2023). An ORF with a q-value < 0.05 in this list (Wacholder, et al. 2023) was considered translated.

Translated ORFs in this study were classified as either “conserved” or “*de novo*” based on previous studies (Carvunis, et al. 2012; Wacholder, et al. 2023; Rich, et al. 2024). We prioritized the conserved class over the *de novo* class in order to be conservative when calling a sequence *de novo*. A translated ORF is considered conserved as long as it was supported by one of the previous studies. These studies define conserved translated ORFs as those found outside the *Saccharomyces* genus (Carvunis, et al. 2012; Rich, et al. 2024) or having high conservation of reading frames (Wacholder, et al. 2023). Subsequently, a translated ORF was considered *de novo* when it was supported by at least one of the studies and is not conserved in the other studies (Carvunis, et al. 2012; Wacholder, et al. 2023; Rich, et al. 2024).

We selected ORFs that passed these translation and evolution criteria from the BarFLEX collection (Douglas, et al. 2012). We then constructed additional strains with the same expression system for ORFs that passed these criteria and combined them with the selected strains from the original BarFLEX collection. These strains formed the BUDY collection after the quality control. The full BUDY collection was used for the screen with four environments in **Figure 2**. A subset of 557 *de novo* translated ORFs in the BUDY collection was selected for the screen in 22 environments. Most of these 557 *de novo* translated ORFs have *de novo* origins supported by more than one study (Carvunis, et al. 2012; Wacholder, et al. 2023; Rich, et al. 2024).

### Yeast strains

#### BUDY strains

Yeast strains and plasmids are listed in **Table S11**. The yeast strains used in the main screens were derived from the BarFLEX collection (Douglas, et al. 2012). For this study, we have expanded the number of ORFs present in the original collection. Using the entry clone collection from the Vidal Lab and the Gateway cloning method (Thermo Fisher, Waltham, MA), we created an additional collection of galactose-inducible plasmids to express the ORFs of interest. These plasmids were then used to transform the background strain used in the BarFLEX collection, creating the expanded version of this collection, BUDY.

Expression plasmids were made by LR recombination between the Entry clones and the Destination plasmid pBY011 using the LR Clonase II enzyme mix (11791020m, Thermo Fisher, Waltham, MA). The recombination reactions were used to transform DH5〈 competent cells (C2987U, New England BioLabs, Ipswich, MA), and positive clones were grown and selected in 2ml of Luria broth media supplemented with 100μg/ml of Ampicillin (J60977, Alfa Aesar).

The NucleoSpin 96 Plasmid kit (740625.4, Macherey Nagel, Allentown, PA) was used to extract the plasmids, and quantification was done using the plate reader SpectraMax M4 (Molecular devices, San Jose, CA).

The expression plasmids were used to transform the yeast strain with the LiAc/PEG/ssDNA transformation protocol (Dunham, et al. 2015) with an adaptation to be performed at a high-throughput scale. The reference strain was constructed using an empty pBY011 plasmid. The background strain was grown in 96-deep well plates (1ml of YPD+G418 per well) and used for transformation performed with the liquid handler EVO 150 (Tecan, Morrisville, NC). Cells were washed in 750μl of water, followed by a washing step in 1ml of LiAc/TE, and finally resuspended in 50μl of LiAc/TE. Transformation was carried out using 5μl of ssDNA carrier DNA 2mg/ml stock (15632011, Invitrogen, Thermo Fisher, Waltham, MA), with 150ng of purified plasmid and 250μl of PEG/LiAc/TE mix. The transformation mix was incubated at 30°C for 15min, followed by 60min at 42°C.

Cells were then pelleted, resuspended in 200μl of SC-URA+glucose+G418 media for a recovery step at 30°C for 3h, prior to being used to seed 3μl drops on SC-URA+glucose+G418 plates. Plates were incubated at 30°C for 2 to 4 days until transformants grew. Agar plates were then used to pin onto 96-well plates containing 100μl of liquid media SC- URA+glucose+G418 with the Singer RoToR (Singer Instruments, Watchet, UK) and then used to make glycerol stocks.

#### Quality control of BUDY

We assessed the quality of the BUDY collection by sequencing the overexpressed ORFs. We prepared PCR templates by colony PCR, which was done by dissolving colonies in lysis buffer (1XTE, 0.25% SDS) and denaturation at 95°C for 10min followed by a quick spin down. The plasmid regions with the ORFs were PCR-amplified with GoTaq polymerase (Promega, Madison, WI) and primers ARC0001 (ACGTTGTAAAACGACGGCCAGT) and ARC0002 (TTTCACACAGGAAACAGCTATGAC). Subsequently, sequencing was performed by GeneWiz (South Plainfield, NJ), and/or Plasmidsaurus (Arcadia, CA). The sequencing results were manually examined. A strain passed the quality control when the protein sequence of the ORFs was identical to the sequences reported in the published translated ORF list (Wacholder, et al. 2023). Strains failing the quality control were filtered out from analyses.

#### Strains for the mechanistic study of *DRD1*

The three strains for the mechanistic study of *DRD1*, DRD1-mNG-OE, control-mNG-OE and parental strains, use a β-estradiol-inducible system. The parental strain used as reference has the following genotype: MATα *CAN1*pr::*TDH3*pr-E2Crimson::KanMx4::*can1*Δ::*STE2*pr- *LEU2 HTA2*-mCherry::Ca*URA3 leu2*Δ0::*ACT1*pr-ZEV3::NAT *lyp1*Δ *ura3*Δ0 *his3*Δ1 *met15*Δ0. The additional overexpression strains contain a construct (Z3EVpr-*YLR112W*-mNG:HPH or Z3EVpr-*YIL0CCW-A*-mNG:HPH) replacing the *his3Δ1* locus and selected by the presence of a Hygromycin cassette and the resistance to the antibiotic Hygromycin B.

### Media for the overexpression screen

The growth media used for this work were based on synthetic complete media lacking uracil (SC-URA), used as selection for the plasmid-based strains, or synthetic complete media (SC) for those strains without plasmids. The carbon source used was glucose (for strain maintenance and any step prior to pre-screening 2 and final screening conditions) or galactose (for pre-screening 2 and final screening conditions).

1.7g of Yeast Nitrogen Base without amino acids and ammonium sulfate (BD Difco), 1g of L-Glutamic acid monosodium acid-free (Sigma, Milwaukee, WI), and 2g of amino acid drop- out mix **(Table S12)** were mixed and dissolved in a final volume of 100ml in water and filter-sterilized, to make a 10X stock solution. For any of the conditions used in the final screenings **(Table S13)** the conditions were adjusted by using a different amino acid drop-out **(Table S12)** to prepare the 10X stock solution (SC-URA, SD-MET or SD-URA+4mg/ml of methionine, “Low MET”), by adjusting the pH of the 10X stock solution before filtration (pH 2.0, pH 3.5, pH 7.0 and pH 9.5) or by adding DMSO or the stressors of interest (rapamycin, tunicamycin, fluconazole, hydroxyurea or H_2_O_2_) at the same time as the antibiotic G418.

Liquid media was prepared by dissolving the 10X stock solution into 1X in 850ml of water, and 50ml of 40% glucose or galactose (Sigma) stock solution.

Agar plates were prepared by dissolving and autoclaving 20g of agar (BD Difco) in 850ml of water. This autoclaved mixture was then brought to 65°C and supplemented with 100ml of the 10X stock solution and 50ml of 40% glucose or galactose stock solution. Each omni plate (PlusPLates PLU-003, Singer Instruments, Somerset, UK) was filled with 50ml of the agar media mixture and dried out for 24-48 hours prior to use.

In both liquid and agar media, the media was supplemented with G418 (G640001.0, RPI, Mt. Prospect, IL) at a final concentration of 200μg/ml, using a stock solution of 100mg/ml.

For the experiments using β-estradiol (E8875-1G, Sigma) as expression inducer, the media of interest, either SC or SD, containing glucose as carbon source were supplemented with β-estradiol 10μM using a stock solution of 2000μM.

### Colony-based high throughput phenotyping

The colony-based high throughput phenotyping was performed the same way unless stated otherwise for three experiments, including: (1) the screen with the full BUDY collection in four environments; (2) the screen with strains of 557 *de novo* translated ORFs from the BUDY collection in 22 environments; and (3) the retest. Colony-based high-throughput screens were conducted at 6,144-colony density. Following the Linear Interpolation Detector (LID) framework for high throughput colony-based screening (Parikh, et al. 2021), the reference strain was grown as grids and occupied one-fourth of the plates for correcting spatial artifacts (Baryshnikova, et al. 2010; Parikh, et al. 2021).

The initial glycerol stock was kept at 384 colony density and sequentially replicated to create the final 6144-colony density using the automated pinning ROTOR + (Singer Instruments). To this aim, each stock plate was defrosted, shaken for 30sec at 1200rpm using the MixTape Eppendorf (Eppendorf, CA) and used to inoculate SC-URA+glucose+G418 agar plates **(Table S13)** at 384-colony density **(Table S14)**. Agar plates were incubated for 48h at 30°C. The set of agar plates was then used to upscale into 1536-colony density, combining the three stock plates with one stock of reference strain in four different combinations **(Table S14)** and incubated for 72h at 30°C with continuous imaging every 4h using the automated imaging system spIMAGER A3 (SCP Robotics, Toronto, Canada). This step, defined as pre-screening 1 (PS1), was performed in SC-URA+glucose agar plates. Next, the PS1 plates were used to replicate into pre-screening 2 (PS2) at 1536-colony density into SC- URA+galactose+G418 agar plates and incubated for 72h at 30°C with continuous imaging every 4h.

Final screening was performed by using PS2 plates to upscale into 6144-colony density **(Table S14)** into the different condition agar plates **(Table S13)**. These agar plates were incubated for >72h at 30°C with continuous imaging every 3h for the screen with the full collection and the retest, and every 4h for the screen with the smaller set over 22 environments. The exceptions are the HighTemp and LowTemp environments, which were performed with separate incubators with manual imaging.

### Spatial correction of colony sizes and growth score calculation

This step was performed the same way unless stated otherwise for three experiments: (1) the screen with the full BUDY collection in four environments; (2) the screen with strains of 557 *de novo* translated ORFs from the BUDY collection in 22 environments; and (3) the retest. Raw images of the colonies were first analyzed using the LID pipeline (Parikh, et al. 2021). The LID first calculated the raw colony sizes from images. Raw colony sizes under 300 pixels were filtered out to avoid mistaking marks of pin pads on the agar surface as colonies. The time points when colonies reached saturation in each environment were used for downstream growth score analyses. The saturation time point was defined as the first time point when the raw colony size increase of the reference was <0.3% compared to the previous time point and had >90% of colonies over 300 pixels. The exceptions were the extremely stressful environments, SD-MET pH7.0 and SD-MET pH9.5, where the last time point of the experiment was used for downstream analysis. Raw images of the selected time points were then manually examined to filter out colony positions affected by experimental artifacts such as small condensation, debris, bubbles in agar surface, etc. Uneven distribution of nutrients in agar plates is a common artifact in colony-based high-throughput screens (Baryshnikova, et al. 2010; Parikh, et al. 2021). LID addresses this issue using the reference colonies, which were grown throughout the plates. Briefly, the reference colonies enable the calculation of a predicted colony size of the reference for every single position in the plates.

The colony sizes of individual overexpression colonies were then normalized by taking their ratios to their respective predicted reference colony sizes. Here we denote the raw colony ratio as *S*. As each translated ORF had multiple colonies descending from a single colony at 384-density plate, outliers were detected and removed for each translated ORF as previously described. Outliers were defined as those with a ratio *S* that was more than three median-adjusted deviations away from the medians (Parikh, et al. 2021).

We then corrected for the row-column effect, another common artifact in colony-based high-throughput screens where colonies in rows or columns closer to edges tend to be bigger (Baryshnikova, et al. 2010). We modified a previously described method (Baryshnikova, et al. 2010) while leveraging the reference colonies grown throughout the agar plates. The general idea is to correct the *S* of each colony based on the row and the column it resides in the plate. The row-column corrected value *S*^′^, which is the growth score, was calculated by dividing *S* of a colony at row i and column j by its row and column correction factors, *r_i_*and *c*_j_ respectively:

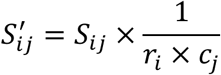

To derive *r_i_* and *c*_j_, the medians of all *S* of the reference colonies in row i and column j, *Mr_i_* and *Mc*_j_, are first calculated to capture the general trends across rows and columns for each plate. Boundary positions, i.e., the four outermost rows (1 to 4 and 61 to 64) and columns (1 to 4 and 93 to 96) near the boundaries of the colony grids were excluded from the calculation. This procedure was done for each plate in each condition, meaning each plate has one vector of *Mr* = [*Mr*_5_, … *Mr_i_*, … *Mr*_60_] and one vector of *Mc* = -*Mc*_5_, … *Mc*_j_, … *Mc*_92_.. A minimum cutoff of eight colonies was set for each row and column to calculate their *Mr_i_* and *Mc*_j_. The approx() function of R was used to estimate *Mr_i_* and *Mc*_j_ for rows or columns where the procedure cannot be applied due to an insufficient number of reference colonies. For example, if *Mr*_11_ is missing in *Mr* = [*Mr*_5_, *Mr*_6_, *Mr*_7_, *Mr*_8_, *Mr*_9_, *Mr*_10_, *Mr*_11_, *Mr*_12_ … ], approx() calculates *Mr*_11_ by linear interpolation using *Mr*_10_ and *Mr*_12_. If a value on either end, *Mr*_5_ for example, is missing, then approx() uses the closest data point, *Mr*_6_, to fill the missing value.

Finally, for each plate, the vectors of row and column correction factors *r* and *c* are calculated by centering *Mr* and *Mc* to 1 with the following equations:

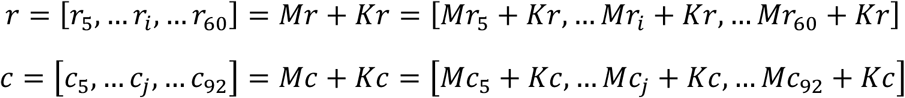

One *Kr* and one *Kc* are decided for each plate with:

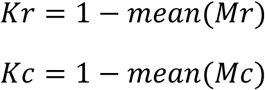

In this way, the means of *r* and *c* are both 1. This keeps a growth score *S*^′^of 1 as the defined point of neutrality after normalization. Then we calculated the means of *S*^′^ for all colonies of each translated ORF. Note that *S*^′^ is the growth score of each colony while the mean of *S*^′^ is the growth score for each translated ORF, we termed them as “colony-level” and “translated ORF-level” data, respectively. The experimental pipeline resulted in 16 replicate colonies in the final 6144-density plates descending from each unique colony in the initial 384-density plates. All strains had one colony in these initial plates, meaning there were 16 colonies per ORF with exceptions: (1) In the screen with full BUDY collection, a small fraction of translated ORFs (168 conserved and 7 *de novo*) have multiple strains. The translated ORF-level data of these strains were derived by first taking the mean of the colony-level data per strain, then taking the mean of the strain-level data; (2) in the retest, each ORF had multiple colonies in the initial plates, resulting in 96 replicate colonies per ORF. For the translated ORF-level data of the reference strain, we applied the same procedure used for the overexpression strains, that is, taking the means for reference colonies descending from each position in the initial 384-density plate because each translated ORF mostly descended from single colonies in the initial 384-density plates. The translated ORF-level growth scores of the reference strain were subsequently used to construct empirical null models in the GXE interaction analysis for the screen with the full BUDY collection and the beneficial/deleterious phenotype calling for the screen in 22 environments.

### Gene-by-environment (GXE) interactions of overexpression phenotype

For the screen with the full BUDY collection in four environments, we calculated interaction scores, *i*, by dividing translated ORF-level growth scores in a stress environment, NaCl or tunicamycin, by those in its corresponding control environment. The interaction scores *i* of the reference strain in each environment pair formed an empirical null distribution to decide whether a specific translated ORF interacted with a stressor. To achieve this, we decided thresholds for positive and negative interactions separately for each stress-nonstress environment pair, such that the empirical FDR was below 0.05. The empirical FDR of each stress-nonstress environment pair was calculated as

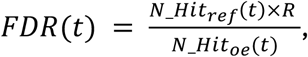

where *t* is the threshold of *i* to call either positive or negative interaction; *N*_*Hit_oe_* and *N*_*Hit_ref_* are the number of the translated ORFs and the reference with *i* more extreme than the threshold *t*; and *R* is the scaling factor, which is the ratio of the number of translated ORFs to the number of references tested in the screen. This procedure was repeated for each combination of stresses (NaCl or tunicamycin), strain types (conserved or *de novo*), and directions of interactions (positive or negative) separately. A significant positive interaction indicates that the stress positively modified the growth score of the translated ORF, and vice versa for a significant negative interaction.

### Phenotype assignment relative to reference strain for the 22-environment screen

To determine whether a translated ORF was beneficial or deleterious when overexpressed relative to the reference strain in each of the 22 environments in the follow-up screen, we applied an approach similar to the GXE analysis. For each environment, the ORF-level growth scores of each overexpression strain were compared to a null distribution formed by the ORF-level growth scores of the reference strain. We calculated empirical FDRs:

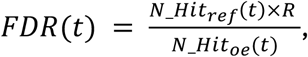

where *t* is the threshold of *i* to call either beneficial or deleterious phenotype relative to the reference strain; *N*_*Hit_oe_* and *N*_*Hit_ref_* are the number of the translated ORFs and the reference with *i* more extreme than the threshold *t*; and *R* is the scaling factor, which is the ratio of the number of translated ORFs to the number of references tested in the screen. This procedure was repeated for each of the 22 environments and directions of phenotypes (beneficial or deleterious) separately. A significant beneficial phenotype means the ORF-level growth score of an ORF is higher than the threshold for beneficial phenotype, and vice versa for a significant deleterious phenotype. Only translated ORFs that were successfully tested in at least 10 environments were included for phenotype assignment to make the environments comparable. Note that we aimed to control the empirical FDRs under 0.05, yet the achieved FDRs are often much more stringent. All environments for beneficial phenotypes and most environments (15/22) for deleterious phenotypes have FDRs approaching 0 **(Table S6).**

### Phenotype assignment relative to reference strain for the retest

In total, 29 translated ORFs were retested over three environments. Around 1800 and 96 colonies were grown for the reference strain and each of the overexpression strains, respectively. The colony-level growth scores of all colonies of each overexpression strain were compared to those of all reference colonies with Wilcoxon Rank Sum tests using the wilcox_test() function of the R package rstatix (Kassambara 2023). BH-adjustment was performed over all 87 comparisons, i.e., 29 translated ORFs over three environments. The phenotype assignment approach in the retest is based on Wilcoxon Rank Sum test instead of the empirical FDR control used in the screen because the goal is to confirm phenotype instead of hit discovery, allowing us to report statistical significance of each ORF- environment pair **(Table S7)**.

### Liquid growth assay and analysis

Liquid assay was performed using the DRD1-mNG-OE, control-mNG-OE, and parental strains **(Table S11)**. Cells were grown in liquid SC + glucose + G418 medium overnight at 30°C with shaking at 220 rpm. On the day of the experiment, cells were diluted 1:10 in fresh media with either 20 μM of β-estradiol (Sigma) for overexpression induction and grew for three hours. Then, cells were pelleted and resuspended with liquid SC + glucose + 100nM rapamycin (R8781-200UL, Sigma) or its vehicle DMSO (D8418-100ML, Sigma) of equal volume **(Table S13)** to reach a final cell density around 0.1 OD_600nm_ and a volume of 200μl per well in a black 96-well plate (655986, Greiner Bio-One, Thermo Fisher). The cell density OD_600nm_ and mNeonGreen fluorescence signal (excitation and emission wavelengths of 506 and 536 nm, respectively) were measured every 20 minutes using a microplate reader BIOTEK SYNERGY H1M (Agilent, Santa Clara, CA). The growth and fluorescent data were analyzed in R. The R package gcplyr (Blazanin 2024) was used to import and analyze per-well the growth parameters, including the growth rate and maximum cell density. Wilcoxon Rank Sum tests and BH correction were done using the R package rstatix (Kassambara 2023).

### Sample preparation for RNA-sequencing

We picked three colonies per strain from glycerol stock streaks and grew them independently downstream. Each colony was considered a unique replicate in the RNA-seq experiment. To maximize the yield of cells, five wells of culture were prepared per strain per environment. We repeated the same liquid growth assay described in the previous section. The cell density at OD_600nm_ and mNeonGreen fluorescence signal were measured every 20 minutes. This ensured that all groups were in the same growth phase (late exponential phase) and that the mNeonGreen signals were present in both overexpression strains.

Once cells were grown for 10 hours and reached the late exponential phase at an OD_600nm_ around 1.4, 5 wells of 200μL culture of each replicate were combined into one 1.7mL tube and spun at 3700 rpm for 5min at 4°C. After spinning, the supernatant was removed, and the cell pellets were rinsed with 500μL of cold 1X phosphate buffer saline (PBS), split into two tubes, and spun again. The supernatant was removed again, and cell pellets were immediately flash frozen and stored at -80°C.

RNA was extracted using Qiagen’s RNeasy Mini kit (74104, Qiagen, Germantown, MD) and following instructions for lysis of <2x10^7^ yeast cells. Specifically, cells were lysed to create spheroplasts by resuspension in 100μL of Buffer Y1 (1M sorbitol, 0.1 M EDTA, pH 7.4) containing 10μL of zymolase by incubating for 10 min at 30°C with gentle shaking. After incubation, 350μL of Buffer RLT prepared as indicated by the kit (10μL of β-mercapto- ethanol to 1mL Buffer RLT) was added to the cell lysate and vortexed vigorously to lyse spheroplasts. This solution was centrifuged for 2min at full speed, and the supernatant was transferred to a clean tube. 350μL of 70% ethanol was added to the lysate and mixed by pipetting before transferring to the RNeasy spin column and spinning for 15sec at max speed, discarding the supernatant. The column was then rinsed once with 700μL of Buffer RW1 and twice with 500μL of Buffer RPE, discarding the supernatant each time. Finally, RNA was collected by adding 30μL of water to the RNeasy spin column and spinning for 1min at max speed. RNA quality was measured by TapeStation (University of Pittsburgh Genomics Core Facility) with RIN scores ranging from 5.5-8.4. Samples were prepared for 3′ end sequencing with Plasmidsaurus (Arcadia, CA) by diluting to 45ng/μL in 65μL of water before adding 1.45μL of 1:5 diluted SIRV-Set 3 spike-in (Lexogen, Greenland, NH).

### RNA-seq data processing and analysis

#### Read mapping

We followed the pipeline described by Plasmidsaurus, accessed 1 May 2026, with minimal modification (Plasmidsaurus 2026). Briefly, raw reads from 18 single-end libraries were quality-filtered using FastP (v0.24.0) (Chen, et al. 2018) with poly-X tail trimming, 3′ quality trimming (Phred Q ≥ 15), and a minimum post-trim read length of 50 bp. The reference genome for mapping consisted of the *S. cerevisiae* R64 genome assembly (Ensembl release 113), sequences of Lexogen SIRV-Set 3, which include both isoform and ERCC spike-in sequences, and the mNeonGreen sequence. Reads were aligned to the reference using STAR (v2.7.11) (Dobin, et al. 2013), with non-canonical splice junctions removed and intron size bounded to 10–3,000 bp (--outFilterIntronMotifs RemoveNoncanonical --alignIntronMin 10 --alignIntronMax 3000). UMI-based deduplication was performed using UMICollapse (v1.1.0) (Liu 2019). Read counts were then quantified over protein-coding exons using featureCounts (Subread v2.1.1) (Liao, et al. 2013, 2014) in strand-specific forward mode with fractional assignment of multi-mapping reads (-s 1 -M -O --fraction). As the read length is short (∼90 bp), reads from the translated ORF-mNeonGreen transgenes were occasionally mapped to the native YLR112W or YIL066W-A loci. We therefore merged the mNeonGreen read counts into either YLR112W or YIL066W-A sequences based on sample sources before differential analysis.

#### Differential expression and Gene by Environment interaction modeling

Differential expression analysis and count normalization were performed in R using DESeq2 (Love, et al. 2014) with a filtering to retain genes with ≥ 10 counts in ≥ 3 samples. The Pearson correlation coefficients between all possible sample pairs using log2-transformed DESeq2- normalized counts of 92 ERCC spike-in sequences were all above 0.97, suggesting no technical bias in any sample during the sequencing. Subsequently, spike-in sequences were removed from downstream analyses. Two complementary models were applied to identify pairwise DEGs and significant genotype-by-environment interaction: (1) a single-factor grouped model for pairwise genotype or environment contrasts, i.e. rapamycin vs DMSO separately for each genotype and pairwise comparison of the three genotypes separately in either rapamycin or DMSO environments, and (2) a full interaction model (∼ genotype + environment + genotype:environment) to detect genotype-by-environment effects via the default Wald tests on interaction coefficients. Log_2_ fold changes were shrunk using the ashr method. DEGs were defined at adjusted p < 0.05 and |log_2_FC| > 0.5 in the pairwise comparison. Genes with dampened rapamycin responses were identified by taking the intersection of the DEGs in the rapamycin vs DMSO comparison of the parental strain and the genes with significant genotype by environment interactions (adjusted p < 0.05). The R function lm() was used to fit a linear model with the rapamycin response in the parental strain as the predictor variable for the rapamycin response in either of the overexpression strains. This allowed the test of statistical significance on whether the slope differed based on which translated ORF was used in the y-axis.

#### Functional and Regulatory Enrichment analyses

Over-representation-based enrichment analysis was performed with the R package gprofiler2 (Kolberg, et al. 2023), querying GO Biological Process (Ashburner, et al. 2000; The Gene Ontology 2026) and TF binding dataset retrieved from TRANSFAC database (Matys, et al. 2006) against a comparison-specific background gene set. An FDR cutoff of 0.05 was used. Electronically inferred annotations were excluded from the analysis, and a maximum term size of 500 genes was applied when querying the GO database. Rrvgo semantic similarity clustering (threshold = 0.7) (Sayols 2023) was used to collapse highly similar GO terms. To improve interpretability, when more than one target motif of a single TF was significantly enriched, the target motif with the highest fold enrichment was used as the representative. "REPRESSOR", one of the significant results in the TF target enrichment analysis, was manually filtered out since it is not descriptive.

## Acknowledgment

We thank the Center for Cancer Systems Biology (CCSB), the Dana-Farber Cancer Institute for providing the entry clones of the BUDY strains. We thank members of the Carvunis lab for helpful feedback. We thank Drs. Vaughn S. Cooper, Dennis Kostka, and Frederick (Fritz) Roth from University of Pittsburgh and Dr. Maitreya Dunham from University of Washington for their constructive feedback. This work was supported by the National Science Foundation grant MCB2144349 awarded to A.-R.C. We used Claude (Anthropic) to assist with proofreading the manuscript text and analysis code; all scientific content, interpretations, and conclusions are our own.

## Data availability

Phenomic data are available as Supplementary materials at Molecular Biology and Evolution online. Fastq files of the RNA-seq experiment are available through the Sequence Read Archive (SRA) under BioProject ID: PRJNA1509105. The scripts are available at: https://github.com/choulin2/yeast_de_novo_ORF_overexpression_202608.

