## Supplementary Figures and Supplementary Table legend for "Environmental stress and phenotypic tradeoff modulate the adaptive potential of novel coding sequences for *de novo* gene birth"

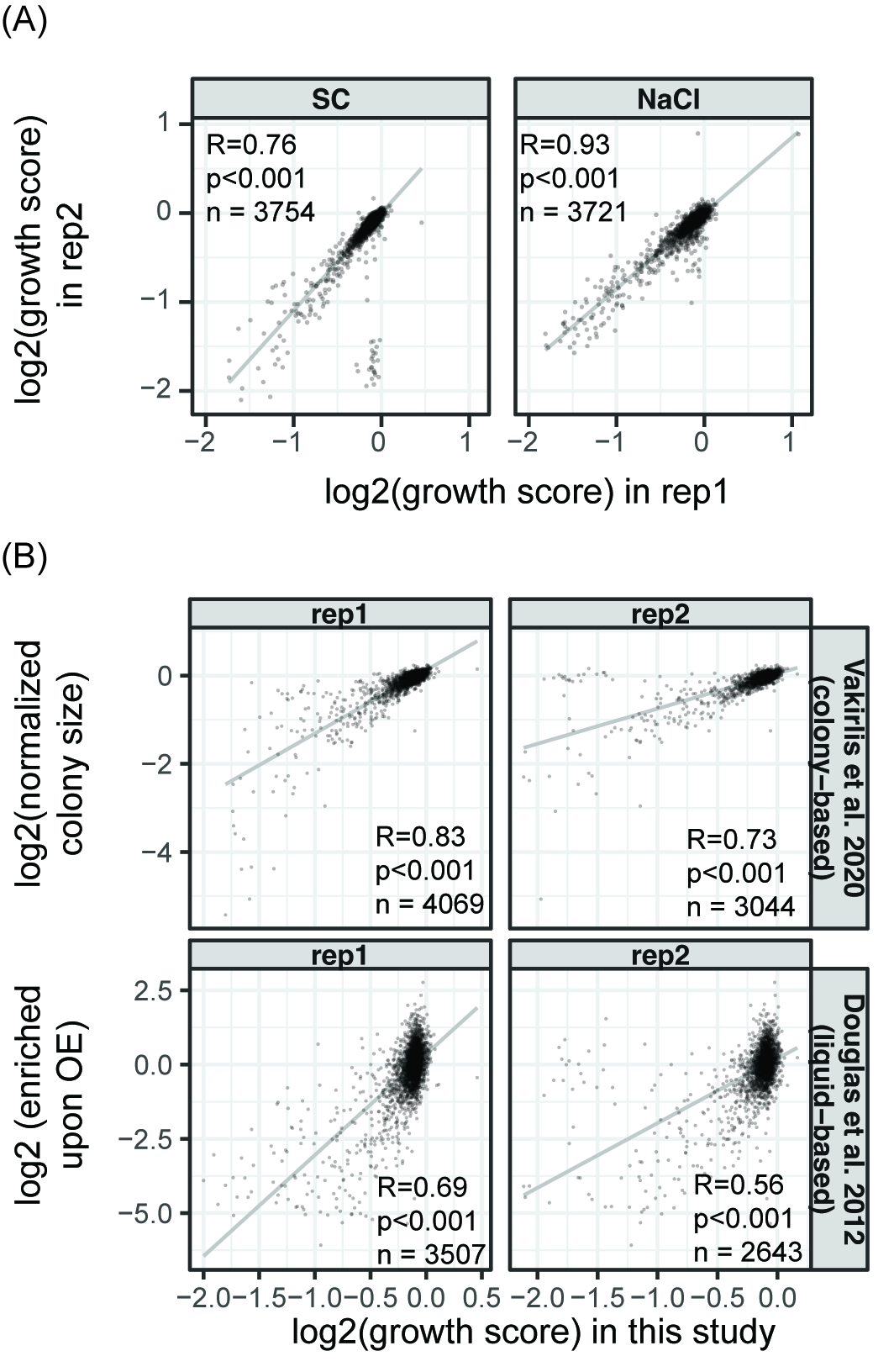

**Supplementary Figure 1.** Comparison of growth scores (A) between replicate experiments of two environments in this study and (B) between the SC environment in this study and previous work that used either colony-based (Vakirlis, et al. 2020) or liquid-based (Douglas, et al. 2012) assays. Pearson’s correlations are shown. Replicate (rep) 1 is used in the downstream analyses. Note that the rep1 vs Vakirlis et al panel in (B) is the same data displayed in Figure 1.
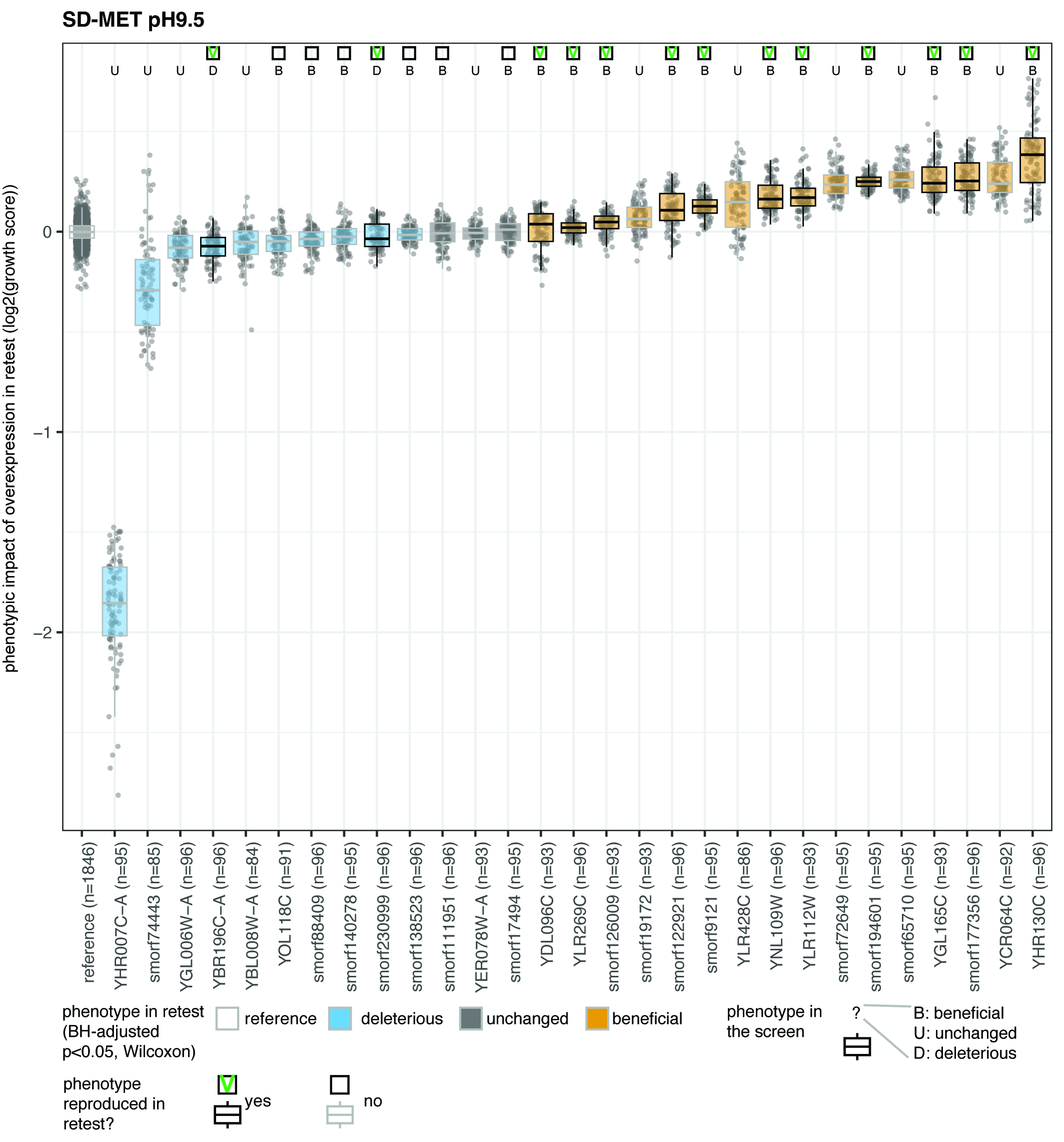

**Supplementary Figure 2.** Retest of 29 overexpression strains in SD-MET pH9.5. Each point represents a colony. Colors filled in the boxes are based on the phenotype in the retest. The phenotypes from the screen are labeled on top. Strains with beneficial or deleterious phenotypes between the screen and the retest are highlighted with darker outline.

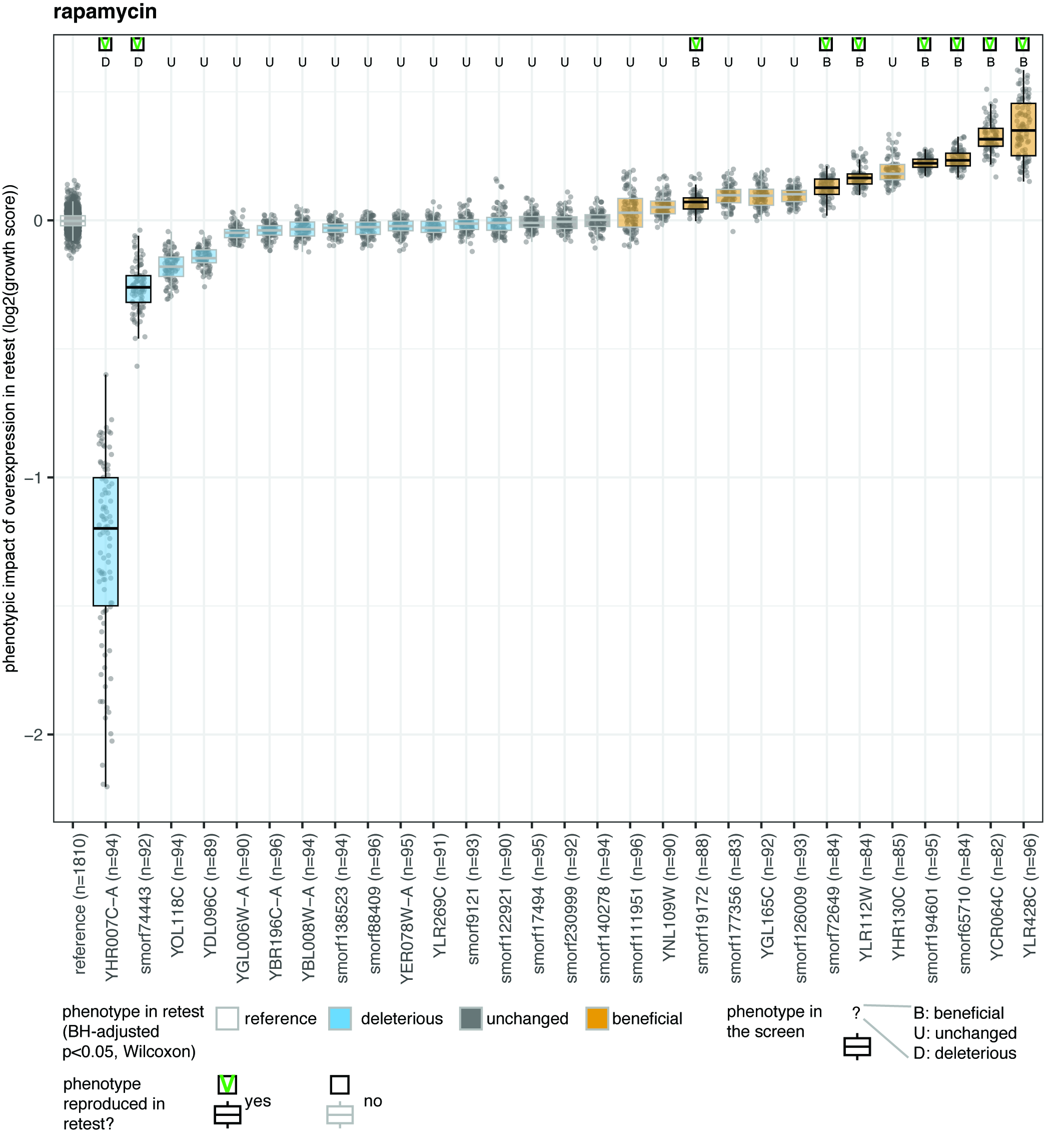

**Supplementary Figure 3.** Retest of 29 overexpression strains in rapamycin stress. Each point represents a colony. Colors filled in the boxes are based on the phenotype in the retest. The phenotypes from the screen are labeled on top. Strains with beneficial or deleterious phenotypes between the screen and the retest are highlighted with darker outline.

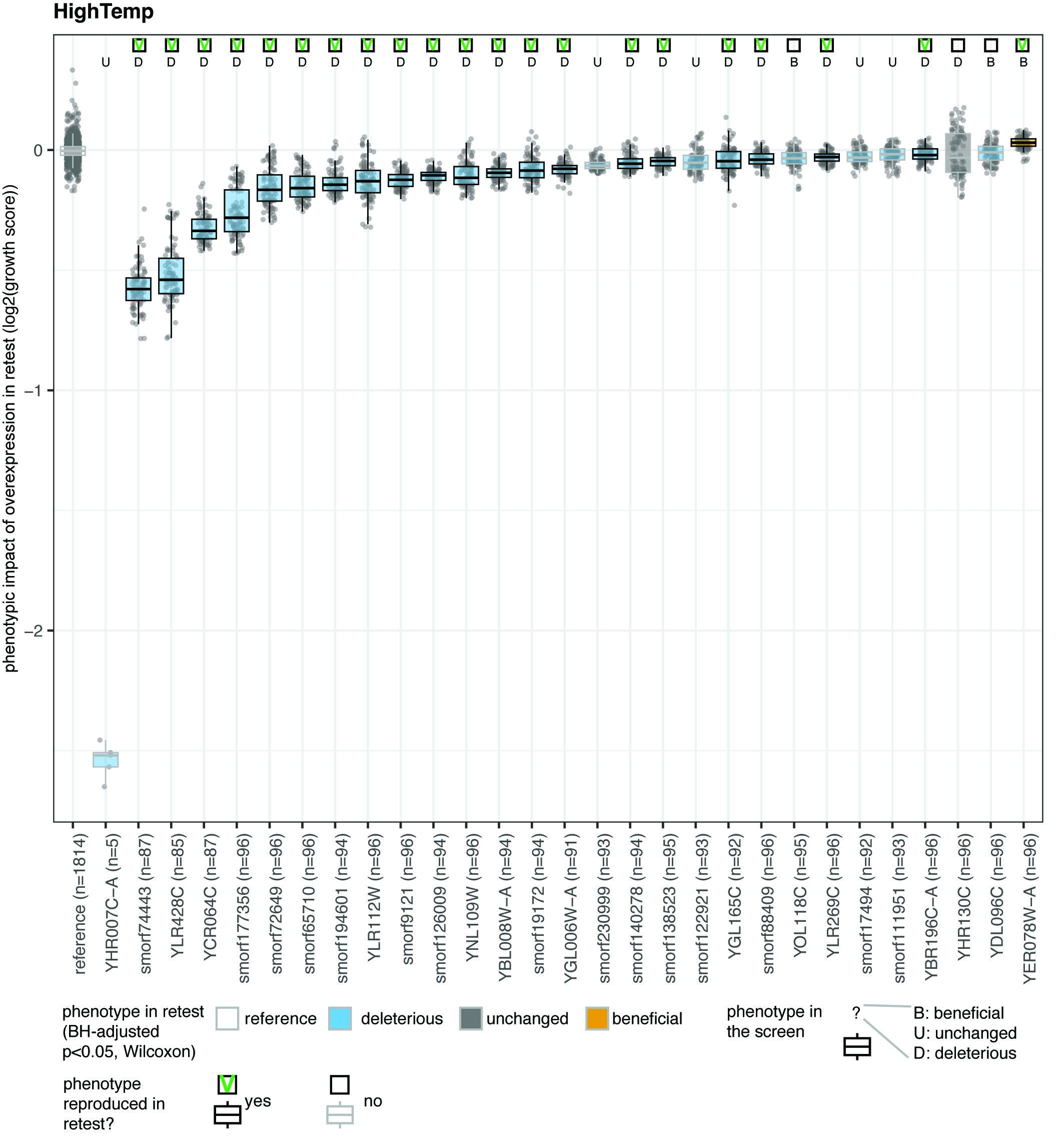

**Supplementary Figure 4.** Retest of 29 overexpression strains in high temperature stress. Each point represents a colony. Colors filled in the boxes are based on the phenotype in the retest. The phenotypes from the screen are labeled on top. Strains with beneficial or deleterious phenotypes between the screen and the retest are highlighted with a darker outline.

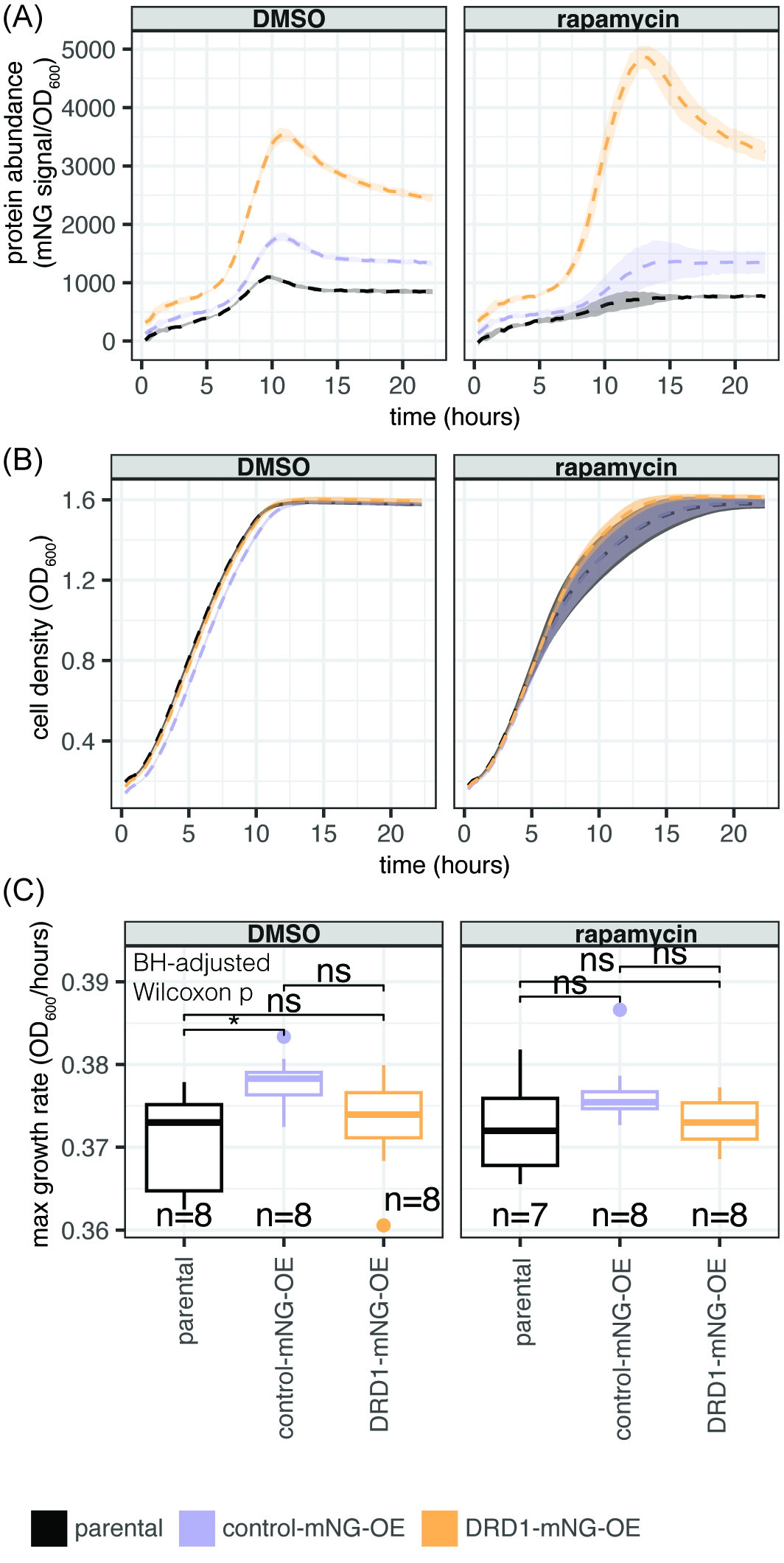

**Supplementary Figure 5.** Liquid growth phenotype and protein abundance of overexpression strains, DRD1-mNG-OE and control-mNG-OE strains. (A) Abundances of Drd1-mNG (in DRD1-mNG-OE strain) and Yil066w-a-mNG (in control-mNG-OE strain), are measured as the fluorescence signal of mNG normalized by the cell density. mNG: mNeonGreen. The results confirm overexpression at the protein level of the two overexpression strains. (B) Growth curves. (C) Comparison of maximum growth rates. Media are based on SC + glucose + β-estradiol.

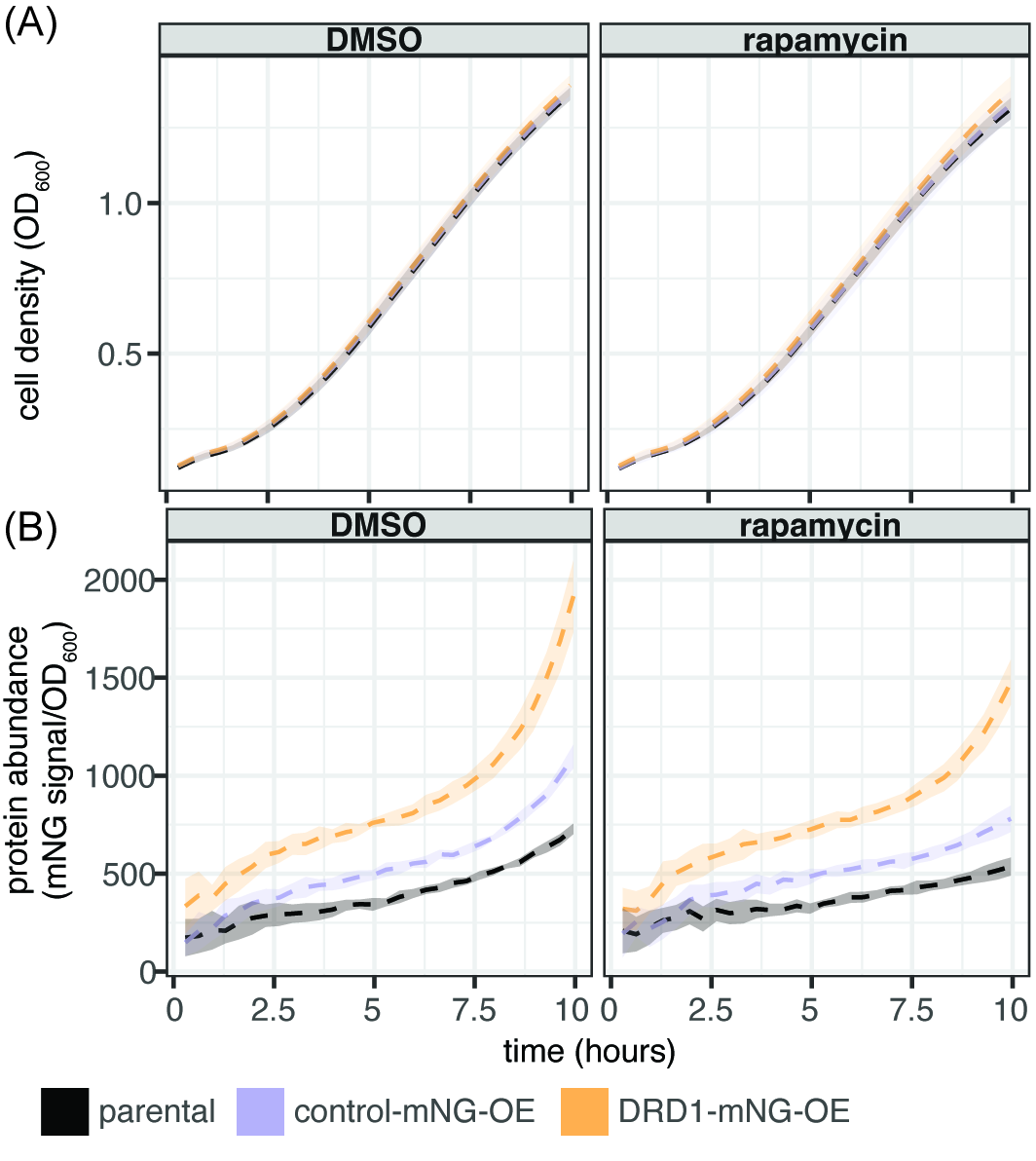

**Supplementary Figure 6**. Liquid growth phenotype and protein abundance of the samples used for the RNAseq. (A) Growth curves. (B) Abundances of Drd1-mNG (in DRD1-mNG-OE strain) and Yil066w-a-mNG (in control-mNG-OE strain), which are measured as the fluorescence signal of mNG normalized by the cell density. mNG: mNeonGreen. Media are based on SC + glucose + β-estradiol.

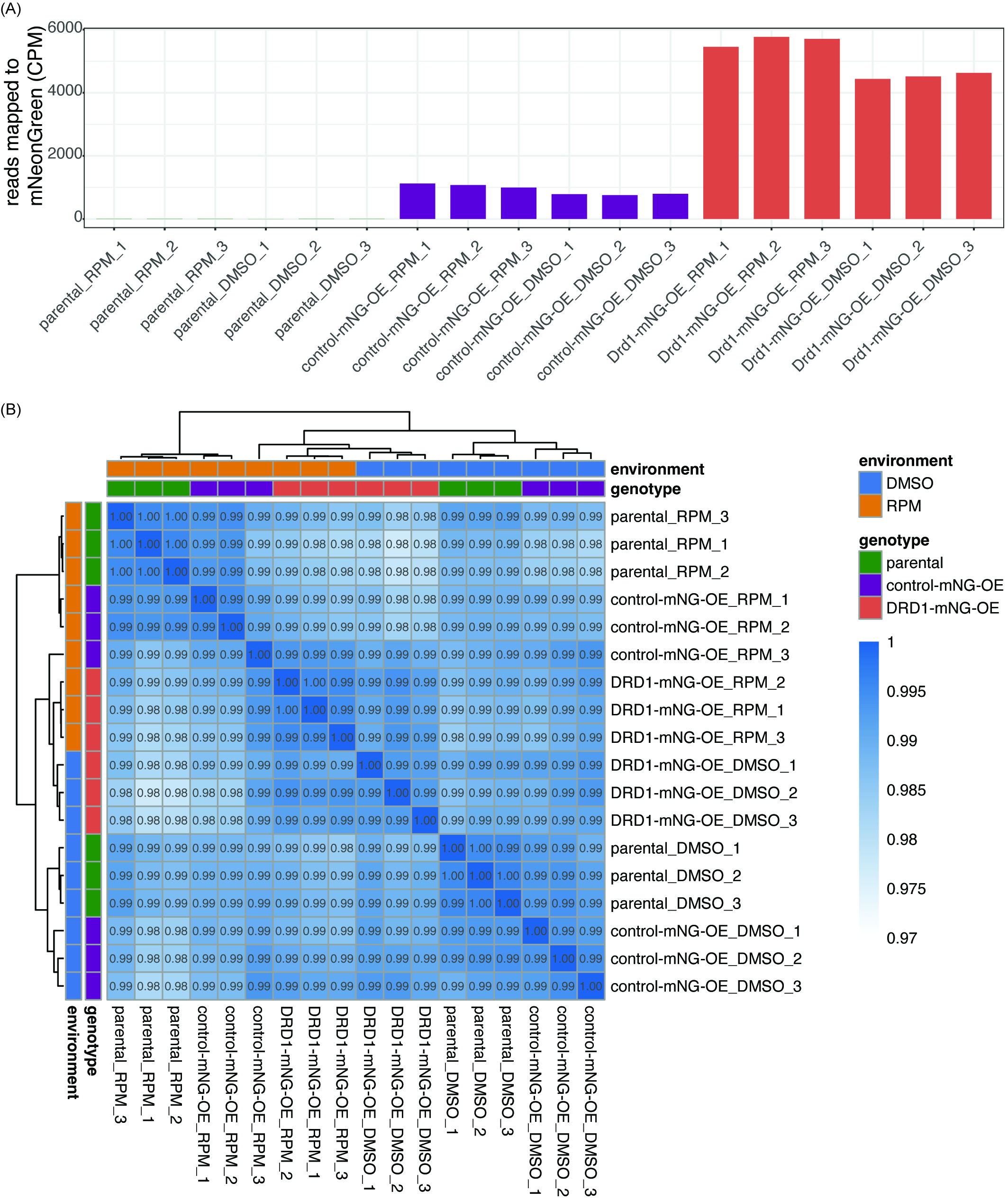

**Supplementary Figure 7.** Confirmation of overexpression at the RNA level and sample correlations. (A) Reads mapped to the mNeonGreen sequence confirm RNA-level overexpression. (B) Spearman’s correlations in expression levels of all genes for all sample pairs. The dendrogram is the average-linkage hierarchical clustering result based on the expression levels after variance-stabilizing transformation in DESeq2. CPM: Counts Per Million. mNG: mNeonGreen.

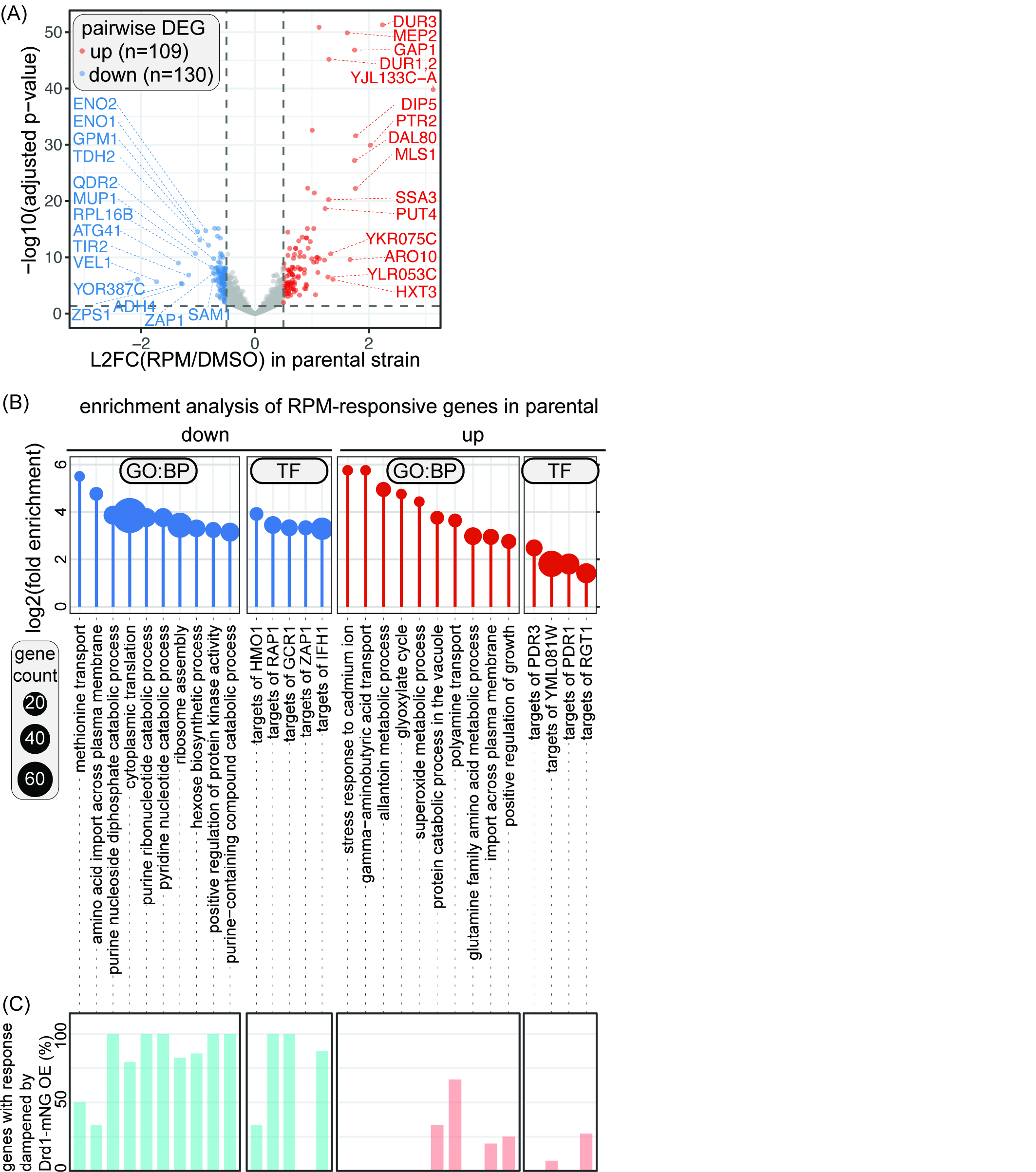

**Supplementary Figure 8.** Transcriptomic response to rapamycin in the parental strain and the dampening effect from Drd1-mNG overexpression. (A) Differentially expressed genes (DEGs) in response to rapamycin in the parental strain. DEGs are defined as those with adjusted p < 0.05 and |L2FC| > 0.5. The 20 most up- and down-regulated DEGs are labeled. (B) GO and TF enrichment analysis of rapamycin-responsive DEGs in the parental strain recapitulates the characteristic growth-suppressing transcriptomic response to rapamycin. (C) Genes that contribute to the enriched terms in the parental strain are dampened by Drd1-mNG overexpression. Bar heights show the proportion of dampened genes in the contributors in each enriched term in (B). RPM: rapamycin. mNG: mNeonGreen. GO: gene ontology. BP: biological process. TF: transcription factor. L2FC: log2 fold change.

**Supplementary tables**

**Table S1.** Strain list of the BUDY collection. Sequence IDs and evolutionary classification from the three references (Carvunis, et al. 2012; Wacholder, et al. 2023; Rich, et al. 2024) are listed and denoted using a suffix. in_original_BarFLEX: whether the strain is from the original BarFLEX collection (Douglas, et al. 2012). “orf_name“ contains the systematic names from SGD or the smorf_id from a reference (Carvunis, et al. 2012) for sequences without systematic names. “seq_verified”: strains with verified sequences of the overexpressed ORFs are marked with “y”. The suffix indicates the company used for the sequencing. Note that a small fraction of ORFs have more than one strain.

**Table S2.** Phenotyping results of the full BUDY collection in four environments. “growth_score_mean” and “growth_score_sd” are the mean and the standard deviation of the growth scores of the colonies of each translated ORF.

**Table S3.** GXE interaction results of the full BUDY collection. “GXE_interaction”: the growth score ratio of a stress environment to its control environment. “env_interaction_FDR0.05”: significant interaction result under empirical FDRs of 0.05.

**Table S4.** List of 557 *de novo* translated ORFs used for the screen over 22 environments.

**Table S5.** Growth scores and phenotype assignment of 557 *de novo* translated ORFs over 22 environments. “growth_score_mean” and “growth_score_sd” are the mean and the standard deviation of the growth scores of the colonies of each translated ORF.

**Table S6.** Empirical false discovery rates of each environment-phenotype direction (beneficial or deleterious) pair of the screen consisting of 22 environments.

**Table S7.** Growth scores in the retest. “growth_score_mean” and “growth_score_sd” are the mean and the standard deviation of the growth scores of the colonies of each translated ORF. “p” is the Wilcoxon Rank Sum test p value. “p.adj” is the BH-adjusted p value. “colony_count_ref” and “colony_count_oe_strain” are the colony counts for the reference strain and the overexpression strain, respectively.

**Table S8.** Descriptive statistics of RNAseq samples.

**Table S9.** Pairwise comparison in RNA-seq. “comparison”: all possible pair-wise comparison in the format of "numerator_vs_denominator". DEG: differentially expressed gene, defined as padj<0.05 and |log2FoldChange| >0.5. “lfcSE”: The standard error estimate for the log2 fold change estimate.

**Table S10.** Genotype-by-environment interaction analysis of the RNA-seq data. “comparison”: interaction between DRD1-mNG overexpression and rapamycin or interaction between control-mNG overexpression and rapamycin. "log2FoldChange" is the log2 fold change of the fold change(rapamycin/DMSO) of an overexpression strain to the fold change(rapamycin/DMSO) of the parental strain. "GXE_interaction" is the statistical significance and direction of the interaction.

**Table S11.** Yeast Strains and Plasmids.

**Table S12.** Amino acid drop-out mixes.

**Table S13.** Media composition.

**Table S14.** Pinning settings.
